# Aberrant gliogenesis and excitation in MEF2C autism patient hiPSC-neurons and cerebral organoids

**DOI:** 10.1101/2020.11.19.387639

**Authors:** Dorit Trudler, Swagata Ghatak, James Parker, Maria Talantova, Titas Grabauskas, Sarah Moore Noveral, Mayu Teranaka, Melissa Luevanos, Nima Dolatabadi, Clare Bakker, Kevin Lopez, Abdullah Sultan, Agnes Chan, Yongwook Choi, Riki Kawaguchi, Nicholas Schork, Pawel Stankiewicz, Ivan Garcia-Bassets, Piotr Kozbial, Michael G. Rosenfeld, Nobuki Nakanishi, Daniel H. Geschwind, Shing Fai Chan, Rajesh Ambasudhan, Stuart A. Lipton

**Author notes:** These authors contributed equally (Dorit Trudler, Swagata Ghatak). Department of Medicine, Indiana University-Purdue University, Indianapolis, IN, USA. Correspondence regarding the manuscript should be addressed to S.A.L.

## Abstract

MEF2C has been shown to be a critical transcription factor for neurodevelopment, whose loss-of-function mutation in humans results in MEF2C haploinsufficiency syndrome (MHS), a severe form of autism spectrum disorder (ASD)/intellectual disability (ID). Here, we use patient hiPSC-derived cerebrocortical neurons and cerebral organoids to characterize MHS deficits. Unexpectedly, we found an aberrant micro-RNA-mediated gliogenesis pathway that contributes to decreased neurogenesis. We also demonstrate network-level hyperexcitability in neurons, as evidenced by excessive synaptic and extrasynaptic activity contributing to excitatory/inhibitory (E/I) imbalance. Notably, the extrasynaptic NMDA receptor antagonist, NitroSynapsin, corrects this aberrant electrical activity associated with abnormal phenotypes. During neurodevelopment, MEF2C regulates many ASD-associated gene networks suggesting that our approach may lead to personalized therapy for multiple forms of ASD.

**One sentence summary:** Autism-like MEF2C^+/-^ patient hiPSC models show miRNA-mediated overproduction of astrocytes and hyperactivity of neurons.

## Main Text

Autism spectrum disorder (ASD) is a group of neurodevelopmental disabilities characterized by impaired social interaction and communication, accompanied by stereotyped behaviors and restricted interests (*1*). It is estimated that 1 in 54 American children develop ASD (*1-3*). Despite extensive research that identified various genetic mutations (e.g., *MECP2, SHANK1-3, NRXN1-3*, etc.) and environmental factors, which are contributory to its etiology (*4, 5*), the molecular mechanisms underlying ASD remain largely unknown, hindering the development of robust diagnostics and effective therapies in this field.

Myocyte enhancer factor 2 (MEF2) comprises a family of transcription factors, which belong to the MADS (MCM1-agamous-deficiens-serum response factor) gene family (*6-10*), with four isoforms that play a role in development, cell differentiation, and organogenesis (*11*). After discovering the MEF2C isoform and showing it is the first family member expressed during brain development (*6, 7*), we and others demonstrated its role in neurogenesis, synaptogenesis, and neuronal survival (*12-15*). Subsequently, murine brain-specific knockout of *MEF2C* at early developmental stages was shown to display electrophysiological, histological, and behavioral deficits reminiscent of Rett syndrome (RTT), a severe form of ASD and intellectual disability (ID) (*12, 16, 17*). In fact, the gene responsible for RTT, *MECP2*, is known to influence the expression level of MEF2C and vice-versa (*18, 19*).

Human genetic studies subsequently established an association between *MEF2C* and human ASD/ID. Morrow et al. identified various MEF2 target genes in their screen for autism genes in human pedigrees with shared ancestry (*20*). Moreover MEF2C activity transcriptionally regulates many other genes that have been linked to ASD (*21, 22*). Recently, human studies from multiple laboratories pointed to defects in *MEF2C* itself in ASD, and *MEF2C* haploinsufficiency was identified as the cause for a severe form of ASD/ID in patients with little or no speech output, stereotypic movements, and learning/memory disorder, often accompanied by epilepsy (*23-26*). These studies estimated that as many as 1.1% of patients with ASD or ID manifest a *MEF2C* abnormality as an autosomal dominant cause.

Disorders associated with *MEF2C* haploinsufficiency have been collectively termed *MEF2C* haploinsufficiency syndrome (MHS) (*26*). Importantly, recent comprehensive transcriptome analyses of autistic brains identified *MEF2C* as one of the frequently dysregulated genes in children with ASD even without *MEF2* mutations *per se* (*21, 22, 27, 28*). For example, a histone-wide association study suggested that the MEF2C locus is dysregulated in ASD patients with other etiologies in addition to MHS (*29*). Collectively, these data argue that finding a successful treatment for the MEF2C haploinsufficiency form of ASD may in fact benefit other forms of ASD as well.

Despite prior animal studies of *MEF2C* heterozygosity to mimic MHS (*16, 30, 31*), mechanisms whereby MEF2C affects neuronal development in the human context of ASD are still poorly understood. Accordingly, here we used MHS patient-derived human induced pluripotent stem cells (hiPSCs) and CRISPR/Cas9 technology for isogenic controls to generate cerebrocortical neurons in 2-dimensional (2D) cultures and 3D cerebral organoids. This approach allowed us to study human neuronal development and function in the presence of control (Ctrl) or patient-relevant mutations in *MEF2C*, and thus to tease out molecular and electrophysiological mechanisms underlying MHS pathophysiology.

### Altered differentiation of MHS hiPSC-derived neurons

To study the molecular mechanisms of MHS in human context, we generated hiPSC lines from dermal fibroblasts of 3 patients (MHS-P1, MHS-P3, MHS-P4) bearing a microdeletion or point mutation in *MEF2C*. As controls, we used gender-matched samples of similar ages consisting of well-characterized hiPSC lines (Ctrl2 and Ctrl3) (*32*). Additionally, using CRISPR/Cas9, we generated an hiPSC line with a microdeletion in *MEF2C* (designated MHS-P2) that could be compared to its cognate isogenic control (Ctrl1; see **table S1)**. In total, we analyzed 4 hiPSC lines with mutations in MEF2C and 3 control lines. All hiPSC lines were characterized and found to be karyotypically normal, positive for pluripotency markers, and capable of differentiating into all germ layers **(fig. S1)**. We differentiated the hiPSCs into cerebrocortical neurons using a dual-SMAD inhibition protocol (*33, 34*). Both MHS patient and Ctrl hiPSCs efficiently generated neural progenitor cells (NPCs), as detected by immunofluorescent labeling with Nestin and SOX2 (**fig. S2A**). The cells were subsequently differentiated into cerebrocortical neurons (hiPSC-neurons) expressing pan-neuronal markers, including β3-tubulin and microtubule associated protein 2 (MAP2). Our differentiation protocol yielded both excitatory and inhibitory neurons, as well as astrocytes (*33*).

To begin to assess the differentiation capacity of the various hiPSC lines, we analyzed neuronal- and astrocyte-selective markers (MAP2 and glial acidic fibrillary protein (GFAP) or S100β, respectively) after 1 month of differentiation from hNPCs. Compared to Ctrl1, we discovered that hNPCs bearing an MHS-related mutations generated more astrocytes, as detected by GFAP and S100β, and fewer MAP2-positive neurons (**Fig. 1, A to C, and fig. S2, B and C**). We also found lower levels of MAP2 protein, as detected by immunoblot (**Fig. 1, D to F**). MAP2 levels were similarly lower in 3-month-old neuronal cultures (**fig. S2, D and E**), indicating that this effect does not appear only at early stages of development, and persists.

**Fig. 1.**
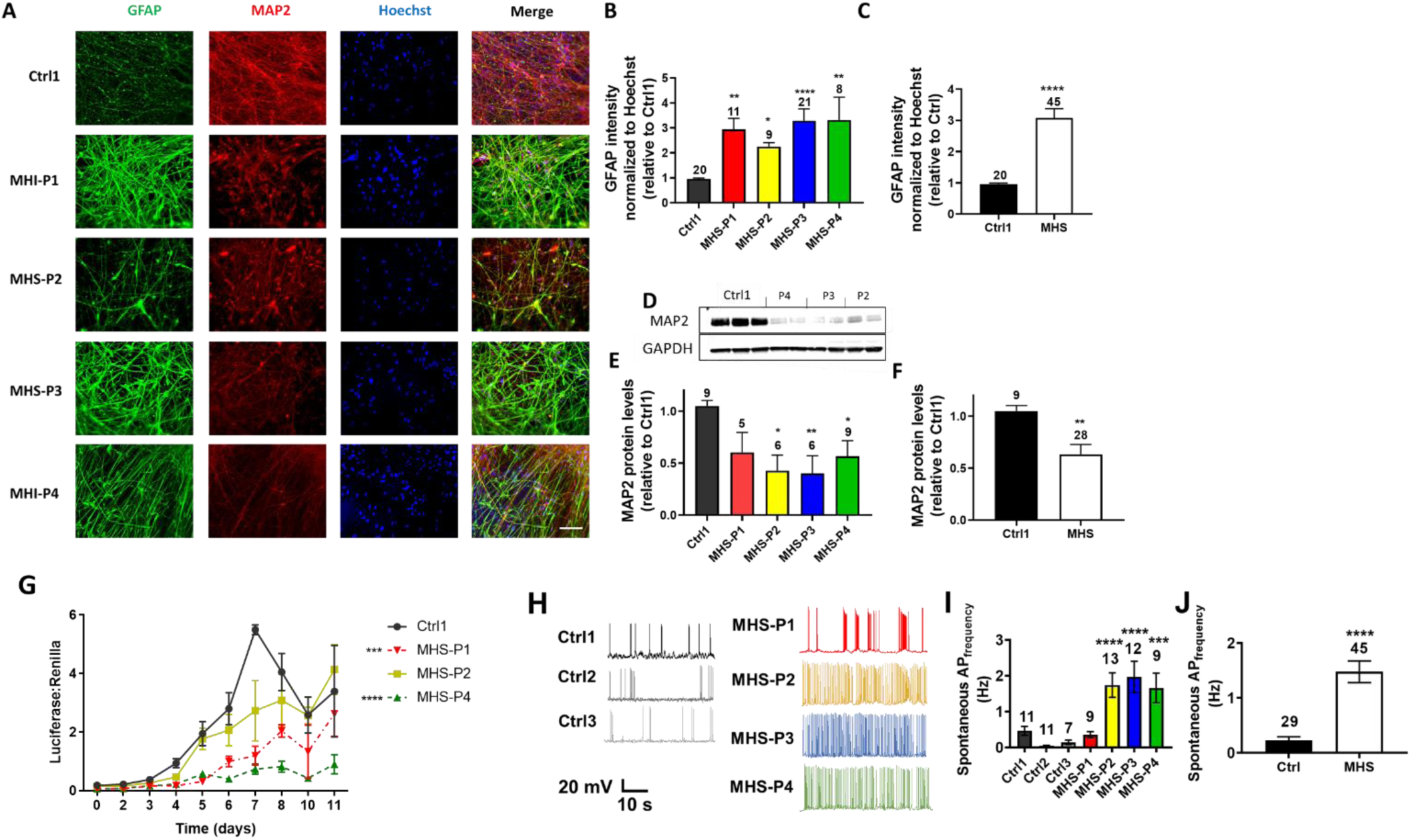
MHS hiPSCs generate more astrocytes and fewer neurons, but with increased spontaneous activity compared to controls. **(A)** Representative images of GFAP, MAP2, Hoechst (to label cell nuclei), and merged images in 4-week-old hiPSC-derived cultures. Patient-relevant mutation-bearing hiPSCs abbreviated MHS-P1 through P4. The genetic background of MHS-P2 is isogenic to Ctrl1. Scale bar, 100 µm. **(B)** Quantification of GFAP fluorescence intensity for each MHS line normalized to Hoechst and relative to Ctrl1 (in this and subsequent figures, other controls yielded similar results). **(C)** Grouped analysis of GFAP expression for Ctrl1 vs. all MHS patients. **(D)** Representative immunoblot of MAP2 expression with GAPDH as loading control. **(E)** Quantification of MAP2 protein expression by densitometry in each MHS line relative to Ctrl1. **(F)** Grouped analysis of MAP2 expression showing a decrease in protein levels in MHS lines vs. Ctrl1. **(G)** MEF2 luciferase reporter gene activity normalized to Renilla at different time points of neuronal differentiation for each MHS line vs. Ctrl1 as a representative control line. **(H)** Recordings of spontaneous action potential (sAP) at resting membrane potential (RMP). **(I)** Quantification of sAP frequency in neurons from each MHS patient (P1-P4) compared to each control (Ctrl1, Ctrl2, Ctrl3). **(J)** Grouped analysis of sAP frequency. Values are mean ± SEM. Sample size listed above bars (number of fields for immunofluorescence panels and number of neurons for recordings, each in ≥3 independent experiments). \**P* < 0.05, \*\**P* < 0.01, \*\*\**P* < 0.001, \*\*\*\**P* < 0.0001 by Student’s t test for pairwise comparisons or ANOVA with Dunnett’s post-hoc test for multiple comparisons.

Additionally, as cells were transitioning from hNPCs to neurons during the first week of terminal differentiation, we performed MEF2-reporter gene assays to begin to assess whether MEF2C participates in the early stages of neuronal differentiation and maturation. We found that the human neurons generated from hiPSCs bearing MHS-relevant mutations (MHS hiPSC-neurons) had lower MEF2 luciferase activity, consistent with haploinsufficiency (**Fig. 1G**). Electrophysiologically, we found that the decrease in MEF2 reporter gene activity correlated with hyperexcitability at both the single neuron and network level that developed within 5 weeks of neuronal differentiation. For example, MHS hiPSC-neurons showed an ∼2-fold increase in frequency of spontaneous action potentials when compared to Ctrl hiPSC-neurons (**Fig. 1, H to J**). We observed this effect for all patient-relevant mutations except MHS-P1, which manifested an increase in spontaneous action potentials at the single neuron level compared to Ctrl2 and Ctrl3 but not Ctrl1 (**Fig. 1, H and I**). This finding may reflect the lack of seizures in this patient compared to the others (**table S1**). Note that evoked action potential frequency and other action potential parameters, including firing threshold, height and half-width, did not show any significant differences between MHS and Ctrl hiPSC-neurons (**fig. S4 and table S2**).

### Transcriptomic effects on MHS hNPCs drive the astrocytic phenotype and limit neurogenesis

To understand mechanistically how MEF2C affects differentiation into neurons, we performed chromatin immunoprecipitation-sequencing (ChIP-seq) analysis on control hNPCs for MEF2C binding sites, using methods we have detailed previously (*35, 36*) (**Fig. 2, A and B and fig. S3**). We found 198 such targets (**table S3**), many of which were enriched for gene ontology (GO) terms related to neurological diseases, cell growth, and proliferation (**Fig. 2A**). Interestingly, five of these MEF2C targets encoded microRNAs (miRNAs) thought to be involved in neurogenesis as opposed to gliogenesis (*37, 38*), so we focused on these (**Fig. 2B**). We first validated the miRNA results using hNPCs that express a constitutively active form of MEF2C (*39-41*) and found that three miRNAs, miR663a, miR663b and miR4273, all of which have binding sites on astrocytic-expressed mRNAs (*42*), manifested higher expression when MEF2C was active (**Fig. 2C**). We then found that patient MHS cells had lower expression of all three of these miRNAs **(Fig. 2D**). When we inhibited the expression of miR663a and miR4273 in Ctrl hNPCs to mimic the effect of haploinsufficiency, we found that the cells recapitulated MHS hiPSC phenotypes, i.e., differentiating mainly into astrocytes and far fewer neurons, as detected by S100β and β3-tubulin (TUJ1), respectively (**Fig. 2, E and F**). These findings are consistent with the notion that suppression of these miRNAs in MHS patient cells contributes to progenitor differentiation into fewer neurons and more astrocytes (**Fig. 2G**).

**Fig. 2.**
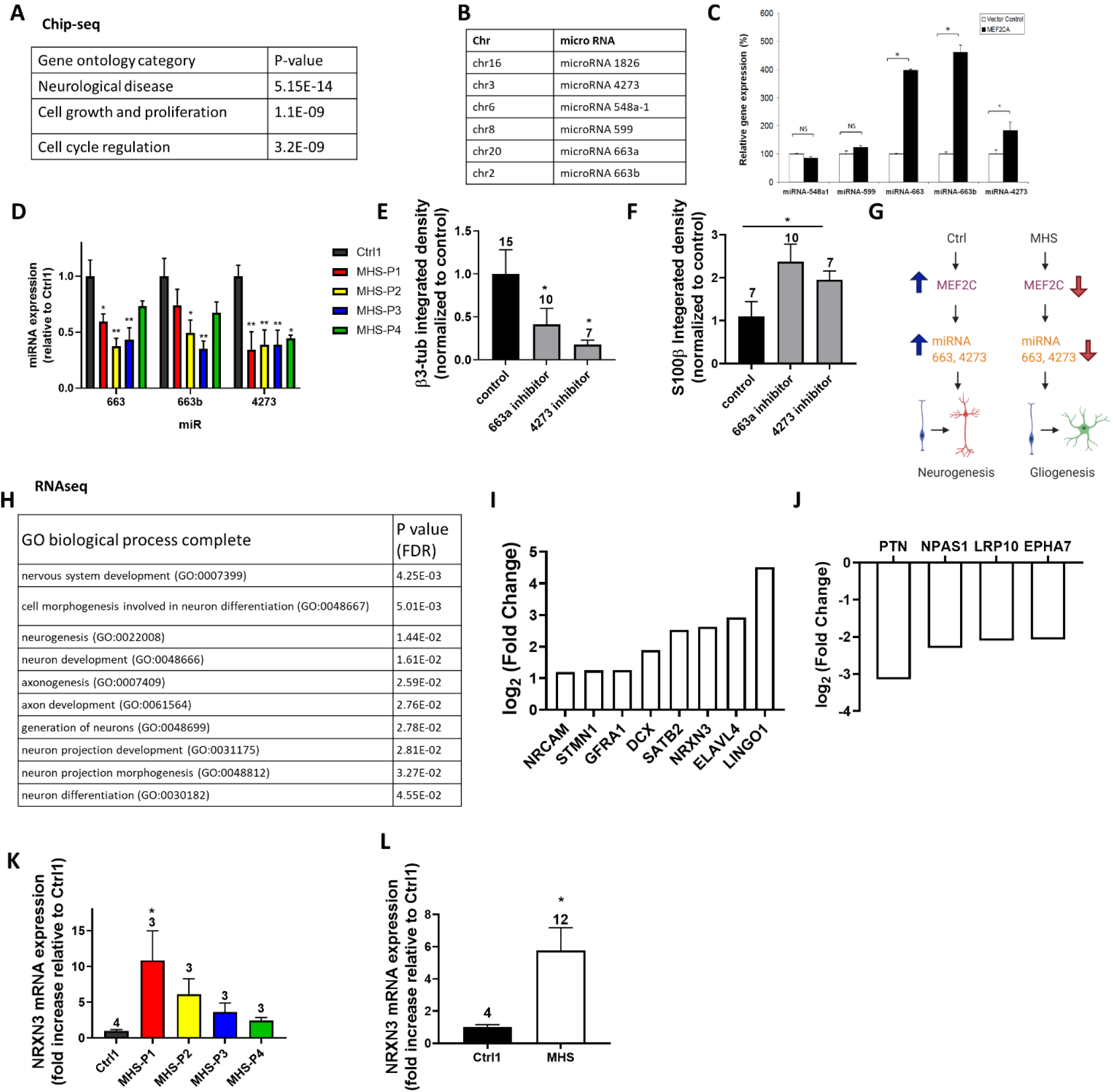
ChIP-seq and RNA-seq analyses show MEF2C effects on hiPSC-derived cell types. **(A)** Top gene ontology (GO) terms for hits found in ChIP-seq analysis of the 198 MEF2C binding targets in hNPCs. **(B)** List of miRNAs identified that contain binding sites for MEF2C. **(C)** Relative gene expression levels of miRNA identified by ChIP-seq in control hNPCs and hNPCs expressing constitutively active MEF2 containing a VP16 transactivation domain (MEF2CA). **(D)** Relative gene expression of miRNA in Ctrl and MHS patient hiPSC-derived cells after 2 weeks in culture. **(E)** β3-tubulin (β3-tub or TUJ11) early neuronal marker expression in Ctrl1 hiPSC-derived cells expressing available miRNA inhibitors compared to non-target control inhibitor after 2 weeks in culture. **(F)** S100β astrocytic marker expression in Ctrl1 cells expressing miRNA inhibitors compared with non-target control inhibitor after 2 weeks in culture. **(G)** Schematic diagram of miRNA effect on neurogenesis and gliogenesis. **(H)** GO terms for differentially expressed neuronal genes by RNA-seq after 5 weeks in culture in MHS patient hiPSC-neurons vs. Ctrl. **(I)** Top differentially-expressed genes from RNA-seq showing higher expression in MHS hiPSC-neurons vs. Ctrl. **(J)** Top differentially expressed genes from RNA-seq showing lower expression in MHS hiPSC-neurons vs. Ctrl. **(K)** NRXN3 mRNA expression in all MHS patients combined vs. Ctrl. Data are mean + SEM. Sample sizes (*n*) are listed above bars from at least 3 independent experiments. \**P* < 0.05, \*\**P* < 0.01 by ANOVA with Dunnett’s post-hoc test for multiple comparisons or by two-tailed Student’s t test for single comparisons.

Next, using methods we have described (*43*), we performed RNA-sequencing (RNA-seq) to detect changes in gene expression between MHS and Ctrl cells after 5 weeks of neuronal differentiation. Previously, we had shown that MEF2C predominantly targets neuronally-restricted gene promoters starting at the hNPC stage (*40*), so we focused our analysis on neuron-associated genes. We identified neuronal genes by their gene ontology (GO) terms (**Fig. 2H**). Among these, several were very highly expressed in neurons compared to astrocytes or other cells types (**Fig. 2, I and J**). Neurexin3 (*NRXN3*) was of particular interest because of its known involvement in neuronal development and function (*44*). Interestingly, neurexins have been associated with ASD, and high or low expression can be detrimental, pointing to a bell-curve effect as found for MEF2C (*45*), with an optimal dose of neurexin critical for normal brain development (*46*). Furthermore, region- and context-specific effects of alterations in NRXN3 have been demonstrated (*44*). Along these lines, we found that NRXN3 was increased in MHS patient hiPSC-neurons compared to Ctrl (**Fig. 2K and L**). Our ChIP-seq and RNA-seq results support the notion that MHS hiPSC-derived cells manifest dysfunctional neurogenesis, increased gliogenesis, and aberrant neuronal differentiation, which we therefore investigated further.

### MHS hiPSC-neurons show enhanced glutamate-mediated currents and decreased GABA-mediated currents

Since both MEF2C and NRXN3 are particularly important for the development of synapses (*12, 17, 30, 31, 44, 47*), we next sought to understand if MHS hiPSC-neurons exhibit differential excitatory and inhibitory activity. We performed patch-clamp electrophysiological recordings on neurons plated on primary mouse astrocytes to enhance synaptic interactions (*48*). Initially, we studied ligand-gated channels. During whole-cell recordings we found the current density elicited by exogenous glutamate application to be 2-4-fold greater in MHS hiPSC-neurons than Ctrl (**Fig. 3, A and B**). In contrast, we found significantly decreased GABA-evoked current density in MHS hiPSC-neurons (**Fig. 3, A and C**). Taken together, this resulted in a robust increase in the glutamate to GABA current ratio (**Fig. 3D**).

**Fig. 3.**
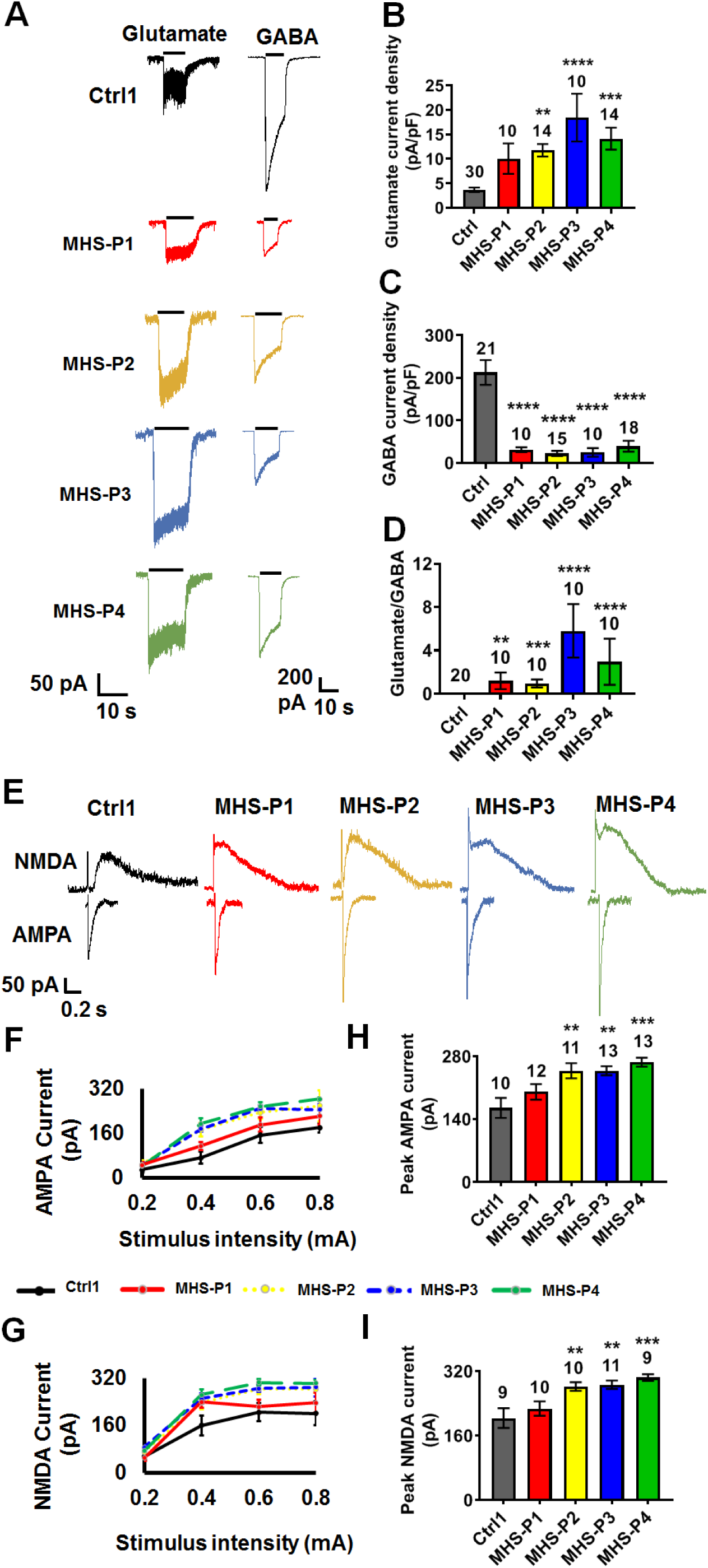
MHS hiPSC-derived cerebrocortical neurons show increased excitation and decreased inhibition. **(A)** Representative traces of glutamate- and GABA-evoked currents (each at 100 µM). **(B** and **C)** Quantification of glutamate and GABA current density. **(D)** Ratio of glutamate to GABA current densities. **(E)** Representative patch-clamp recordings of evoked AMPAR-EPSCs at holding potential (V_h_ = −70 mV) and NMDAR-EPSCs (V_h_ = +60 mV). **(F** and **G)** Input-output curves of evoked AMPAR-EPSCs and NMDAR-EPSCs. **(H** and **I)** Quantification of peak current amplitude of evoked AMPAR-EPSCs and NMDAR-EPSCs. Data are mean ± SEM. Number of neuronal recordings (*n*) listed above bars from at least 4 experiments in each case. \**P* < 0.05, \*\**P* < 0.01; \*\*\**P* < 0.001, \*\*\*\**P* < 0.0001 by ANOVA with Dunnett’s post-hoc test for multiple comparisons (see Methods).

Next, we monitored evoked excitatory synaptic activity. Compared to control hiPSC-neurons, we found a statistically significant increase for MHS-P2-4 (with a trend for MHS-P1) in the NMDA receptor-mediated component of excitatory postsynaptic currents (NMDAR-EPSCs), monitored at +60 mV to ensure removal of Mg^2+^ block by depolarization. There was also an increase in the AMPA receptor-mediated component (AMPAR-EPSCs), measured at -70 mV to ensure separation from the NMDAR-mediated component by blocking NMDAR-operated channels via hyperpolarization-induced Mg^2+^ block (**Fig. 3, E to I**). Too few evoked inhibitory synaptic currents were observed under our conditions to quantify, possibly because of prior findings from the Fishell group showing that MEF2C also plays a role in generating parvalbumin (PV)-positive GABAergic interneurons and as our group has reported in MHS-model mice *(31, 49)*. Other electrophysiological parameters, including resting membrane potential, cell capacitance, and sodium- and potassium-current density, were similar among MHS and Ctrl hiPSC-neurons (**table S2**).

### MHS hiPSC-neurons exhibit disrupted spontaneous synaptic transmission

To further examine differences between excitatory and inhibitory signaling, we next focused on spontaneous synaptic activity. We observed a statistically significant increase in the frequency of spontaneous (s)EPSCs in all MHS hiPSC-neurons except MHS-P1, which manifested a trend in this direction (**fig. S5, A and C**). Additionally, sEPSCs recorded from MHS-P3 and -4 exhibited increased amplitude compared to Ctrl hiPSC-neurons (**fig. S5, A and B**). Concerning spontaneous miniature EPSCs (mEPSCs), we found an increase in frequency with no change in amplitude for most MHS hiPSC-neurons (**Fig. 4, A to C**). MHS-P3, however, displayed a small increase in mEPSC amplitude. In contrast to increased excitatory synaptic transmission, we observed a significant decrease in the frequency of spontaneous and miniature inhibitory postsynaptic currents (sIPSCs and mIPSCs) in MHS hiPSC-neurons compared to Ctrl, without a significant change in amplitude (**Fig. 4, D to F; fig. S5, D to F**). Notably, we had previously reported similar findings in a *MEF2C* heterozygous mouse model of MHS that displays increased excitatory synaptic transmission and decreased inhibitory synaptic transmission (*31*).

**Fig. 4.**
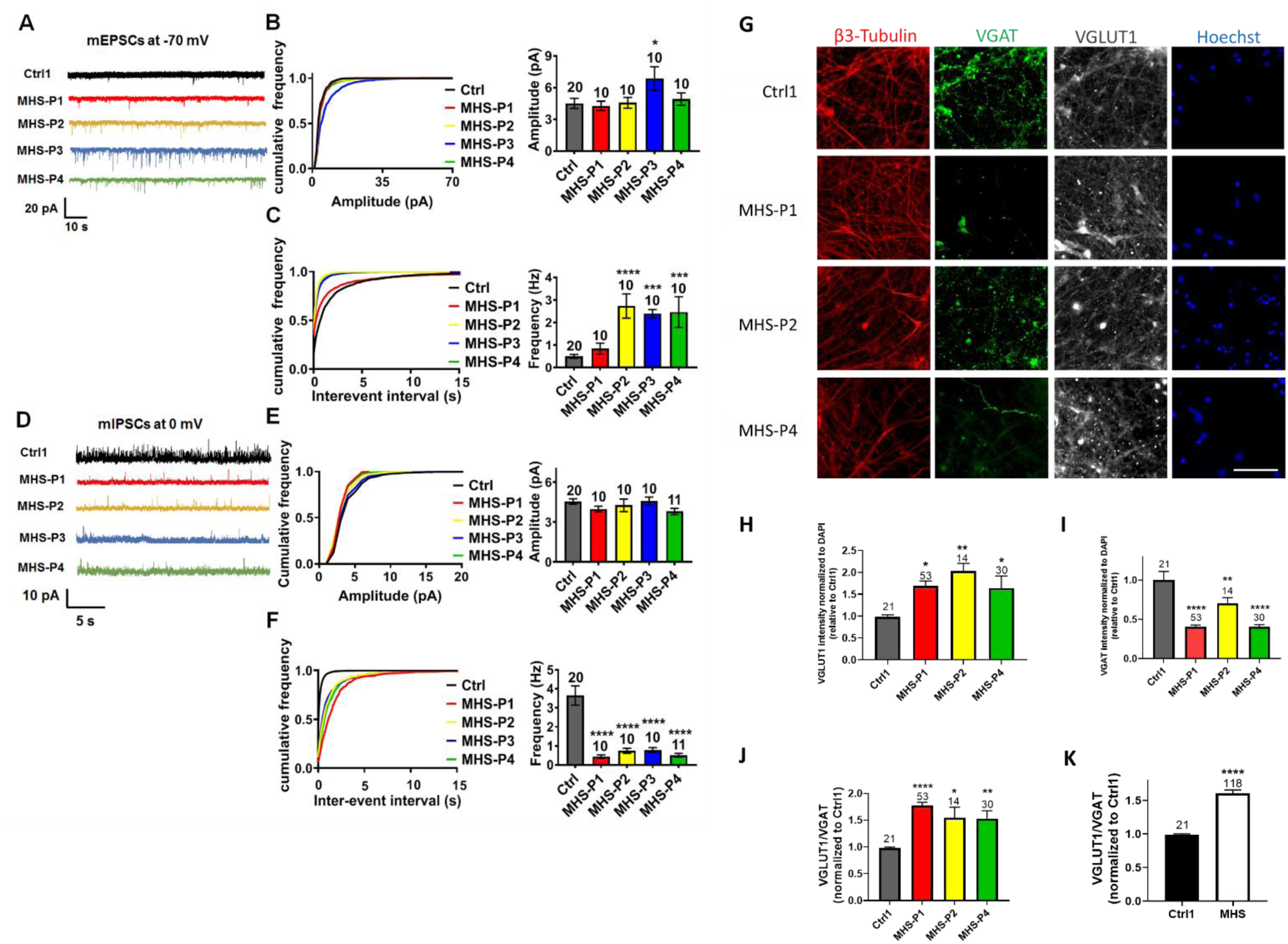
MHS hiPSC-derived cerebrocortical neurons exhibit disrupted synaptic transmission. **(A)** Representative mEPSCs recorded at -70 mV in the presence of 1 µM TTX from Ctrl1 and MHS hiPSC-neurons in culture for 5 weeks. **(B** and **C)** Cumulative probability and quantification of mean mEPSC amplitude and interevent interval (inversely related to frequency). Cumulative probability of MHS mEPSC interevent interval was significantly decreased compared to Ctrl (*P* < 0.0001 by Kolmogorov-Smirnov test). **(D)** Representative mIPSCs recorded at 0 mV. **(E** and **F)** Cumulative probability and quantification of mean mIPSC amplitude and interevent interval. Cumulative probability of MHS mIPSC interevent interval was significantly increased compared to Ctrl (*P* < 0.0001 by Kolmogorov-Smirnov test). **(G)** Representative images of β3-tubulin, VGLUT1, VGAT, and Hoechst STOP in Ctrl1 and MHS hiPSC-neurons. Scale bar, 100 µm. **(H)** Quantification of VGLUT1 in various MHS hiPSC-neurons compared to Ctrl1. **(I)** Quantification of VGAT in various MHS patient neurons compared to Ctrl1. **(J)** Quantification of the VGLUT1/VGAT ratio in various MHS patients. **(K)** Summary of VGLUT1/VGAT in Ctrl1 vs. MHS as a group. Data are mean + SEM. Number of neuronal recordings or imaged fields (*n*) listed above bars from at least 4 experiments in each case. \**P* < 0.05, \*\**P* < 0.01, \*\*\**P* < 0.001, \*\*\*\**P* < 0.0001 by ANOVA with Dunnett’s post-hoc test for multiple comparisons (see Methods).

The increase in mEPSC and decrease in mIPSC frequencies may reflect alterations in presynaptic mechanisms, such as quantal release, or a change in excitatory and inhibitory synapse numbers, or both types of effects. To begin to distinguish between these possibilities, we performed a series of immunocytochemical experiments to monitor synaptic properties. Initially, we measured the total number of excitatory synapses by counting co-localized punctae of presynaptic synapsin I and postsynaptic PSD-95, and found that MHS hiPSC-neurons displayed a moderate, statistically significant decrease in the total number of synapses compared to Ctrl (**fig. S6**). Next, we monitored presynaptic vesicle proteins and observed an increase in vesicular glutamate transporter (VGLUT1) staining but a decrease in vesicular GABA transporter (VGAT), resulting in an increase in the ratio of VGLUT1 to VGAT in MHS hiPSC-neurons compared to Ctrl (**Fig. 4, G to K**). These changes are similar to those we had observed in *MEF2C* heterozygous mice (*31*) and consistent with a presynaptic effect on neurotransmission. The findings on excitatory presynaptic terminals may reflect in part the known action of NRXN3 on VLGUT1 expression and clustering (*44, 46, 47*). Collectively, our results are consistent with the notion that despite a decrease in the total number of excitatory synapses, MHS hiPSC-neurons may exhibit enhanced presynaptic release, resulting in an overall increase in excitatory neurotransmission. Moreover, in addition to its role in neurogenesis and synaptogenesis during development (*12*), MEF2C is also known to play a critical role in excitatory synaptic pruning once synapses have been formed (*15, 50-52*). Hence, the relative deficit in MEF2C transcriptional activity in MHS hiPSC-neurons compared to Ctrl could also contribute to the increase in mEPSC frequency as a result of decreased excitatory synaptic pruning of MHS hiPSC-neurons as they mature. In contrast to excitatory neurotransmission, we found that inhibitory transmission is relatively depressed by both electrophysiological and histological parameters in MHS hiPSC-neurons. Taken together, the increase in excitation and decrease in inhibition that we observed in MHS hiPSC-neurons may lead to hyperexcitability in the neural network, thus contributing to ASD-like pathophysiology in MHS patients. Therefore, in the next series of experiments we investigated the properties of neural network activity in MHS vs. Ctrl using hiPSC-derived 2D cultures and 3D cerebral organoids.

### MHS hiPSC-neurons exhibit enhanced neural network activity, reflected in spontaneous calcium transients and multi-electrode array recordings, that can be normalized by the extrasynaptic NMDAR antagonist NitroSynapsin

Initially, to investigate neural network activity in a population of neurons simultaneously, we monitored Ca^2+^ transients in 2D cultures. MHS hiPSC-neuronal cell bodies displayed an increase in spontaneous Ca^2+^ transient frequency compared to Ctrl, as monitored with the single-wavelength fluorescent Ca^2 +^ indicator Fluo-4 (**Fig. 5, A and B and movies S1 and S2**). Treatment with the NMDAR antagonist NitroSynapsin normalized the aberrantly high intracellular Ca^2+^ levels and Ca^2+^ transient frequency in MHS hiPSC-neurons toward Ctrl values **(Fig. 5, C to E and movie S3**). The concentration of drug used in these experiments was determined from our previously published dose-response curves that found 5-10 μM NitroSynapsin to be maximally efficacious and also attainable in vivo (*31, 32, 53*). Notably, we had previously reported that NitroSynapsin ameliorates E/I imbalance in MHS-model mice, normalizing their aberrant synaptic activity, histological deficits, and ASD-like behavioral phenotypes (*31*).

**Fig. 5.**
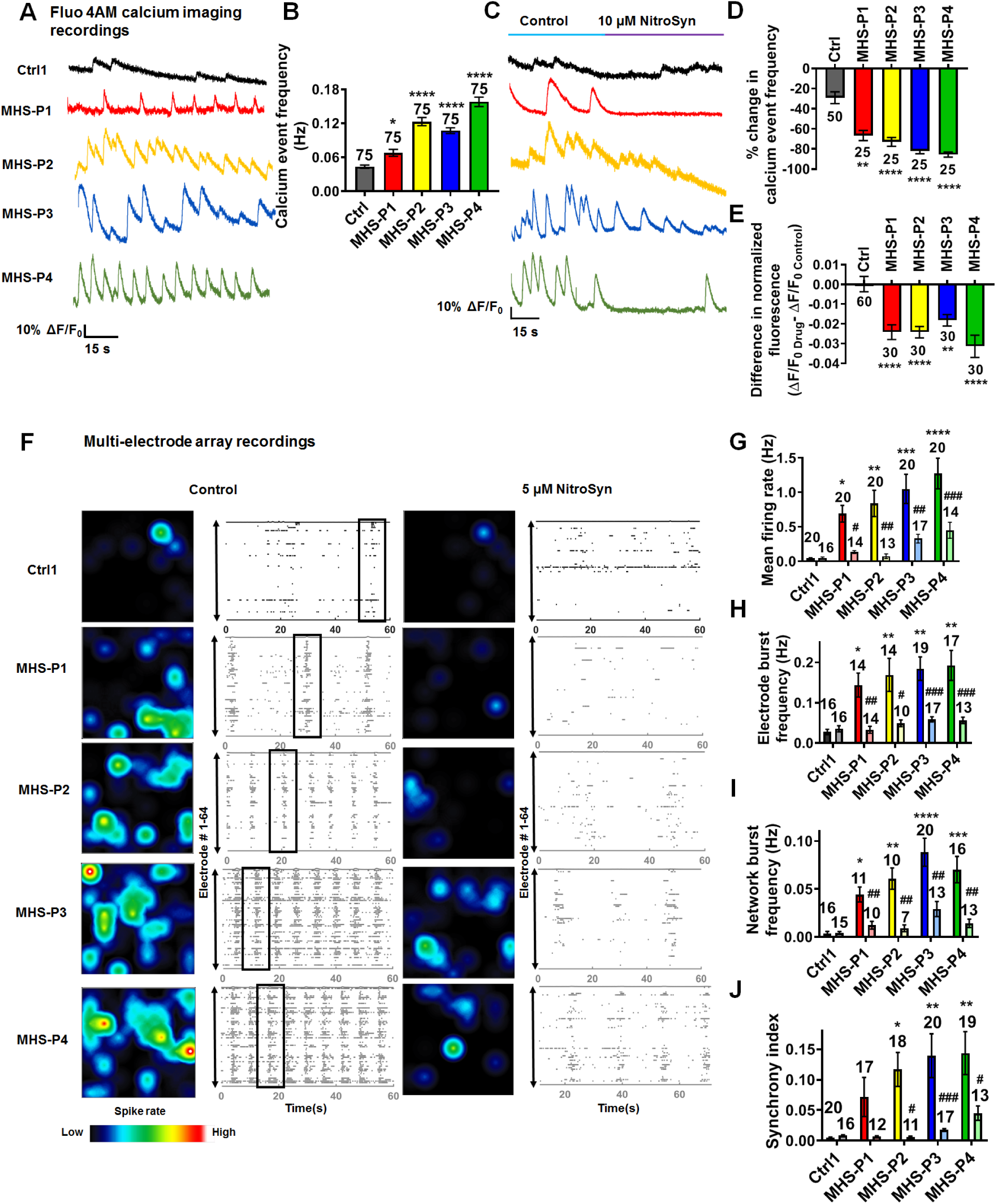
NitroSynapsin normalizes spontaneous calcium transients and neural network activity in MHS hiPSC-derived cerebrocortical neurons. **(A)** Spontaneous neuronal calcium transients recorded from individual Ctrl1 and MHS hiPSC-neurons loaded with Fluo-4 AM. **(B)** Quantification of Ca^2+^ transient frequency for events with rise times <200 ms. **(C)** Representative calcium traces showing decrease in spontaneous calcium transient frequency after application of 10 µM NitroSynapsin (NitroSyn). **(D)** Quantification of calcium transient frequency before and after application of NitroSynapsin. **(E)** Quantification of difference in normalized fluorescence (ΔF/F_0Drug_ -ΔF/F_0Control_) as area under the curve (AUC) in response to NitroSynapsin. **(F)** Representative heat maps and raster plots of MEA recordings from Ctrl and MHS hiPSC neurons before (w/o) and after treatment with 5 µM NitroSynapsin. Boxes outline examples of network bursts. **(G-J)** Quantification of MEA recordings by mean firing rate, electrode burst frequency (representing bursting of individual neurons), network burst frequency (representing bursting of the entire neural network), and synchronous firing. Ctrl represents pooled data for Ctrl1 and Ctrl 2. Data are mean ± SEM. Sample size listed above bars represents number of cells analyzed in 5-10 independent experiments. *^,#^*P* < 0.05, **^,##^*P* < 0.01, ***^,###^*P* < 0.001, ****^,####^*P* < 0.0001 by ANOVA for comparison with Ctrl value (*), or within a group (^#^) for without treatment (w/o) vs. NitroSynapsin treatment; for panel D, comparison was made by non-parametric Kruskal-Wallis test (see Methods).

Mechanistically, we and others have shown that compounds in the aminoadamantane class, such as memantine and the more efficacious NitroSynapsin, preferentially block extrasynaptic (e)NMDARs over synaptic receptors at the concentrations utilized here (*32, 53-55*). Therefore, we next studied eNMDAR-mediated currents in Ctrl and MHS hiPSC-neurons. eNMDAR-associated responses were isolated pharmacologically using a previously published protocol (*32, 56*). Under these conditions, MHS hiPSC-neurons manifested an increase in basal eNMDAR-mediated intracellular calcium responses compared to Ctrl (**fig. S7, A to C**), apparently due to an increase in basal extracellular glutamate levels in the MHS cultures (*32, 56*). Critically, we found that NitroSynapsin could significantly decrease the eNMDAR-mediated calcium responses in MHS cultures (**fig. S7, A and B**). Moreover, treatment with TTX abrogated the increase in basal eNMDAR-mediated extrasynaptic currents in MHS cultures and thus the effect of NitroSynapsin, consistent with the notion that excessive excitatory synaptic activity (as shown in **Figs. 3 and 4**) caused glutamate spillover and consequently an increase in basal eNMDAR-mediated current (**fig. S7C**) (*31, 32, 57*). Additionally, (2R)-amino-5-phosphonovalerate (APV), a competitive NMDAR antagonist, decreased the extrasynaptic currents in MHS cultures, corroborating the notion that the currents inhibited by NitroSynapsin were indeed due to aberrantly active eNMDARs (**fig. S7D**).

Next, we confirmed the presence of aberrantly increased basal eNMDAR-mediated currents in MHS cultures by patch-clamp electrophysiological recordings. We found that MHS but not Ctrl hiPSC-neurons manifested basal eNMDAR-mediated current after pharmacological isolation of this current (*58*), and NitroSynapsin inhibited the current (**fig. S7E**). eNMDAR-mediated currents in spatially-compact neurons would be expected to depolarize their presynaptic terminals and thus alter excitatory synaptic activity. Along these lines, prior studies have shown that aberrant eNMDAR activity can increase mEPSC frequency, probably by increasing presynaptic release of glutamate, as seen here (*59, 60*). Thus, increased eNMDAR currents can lead to a type of feed-forward hyperexcitability, as observed in MHS hiPSC-neurons, and normalizing eNMDAR activity, e.g., with NitroSynapsin, could be critical in rebalancing the neural circuit.

To further explore this effect of excessive eNMDAR activity due to MEF2C haploinsufficiency on neural network activity, we employed multielectrode array (MEA) recordings in our Ctrl and MHS cultures. Similar to our measurements of Ca^2+^ transient activity, in MHS cultures compared to Ctrl we found an increase in mean firing rate, electrode burst frequency (reflecting primarily single neuron action potentials), network burst frequency (reflecting action potentials of multiple neurons), and synchronicity of firing (**Fig. 5, F to J**). Notably, treatment with NitroSynapsin, to block excessive eNMDAR-mediated currents, significantly decreased all of these parameters of aberrant hypersynchronous firing in MHS hiPSC-neuronal cultures while not affecting Ctrl cultures (**Fig. 5, F to J**).

### NitroSynapsin corrects hypersynchronous excessive activity in MHS cerebral organoids

To further validate our results in a more complex system, we next used hiPSC-derived cerebral organoids (*61*), a model system that allows us to study neuronal circuits in 3D, and better reflects in vivo microcircuits (**fig. S8A**). We characterized our MHS and Ctrl cerebral organoids histologically at various time points to confirm the presence of neurons and astrocytes. For example, we found that by 2-months in culture, MHS cerebral organoids compared to Ctrl were smaller in size (**fig. S8, A and B**), similar to human MHS patient brains in both children and adults (*22, 24*). We found that MHS cerebral organoids recapitulated the 2D culture finding of a decreased proportion of neurons and neuropil compared to Ctrl, as judged by quantitative confocal immunofluorescence for β3-tubulin corrected for cell number (**fig. S8, C to E**). We also observed a significant increase in GFAP expression by quantitative confocal immunostaining and by quantitative (q)PCR, but no difference in astrocyte-specific S100β staining (**fig. S8, C, F to K**). Since GFAP labels both radial glial cells/neural precursors and astrocytes but S100β only astrocytes, this finding may be attributed to the developmental stage of these organoids, at which time point neural progenitors were prominent and gliogenesis was just beginning (*62-64*). MHS cerebral organoids also showed an increase in Nestin-positive hNPCs, indicating an abnormal build-up of early neural progenitors compared to Ctrl (**fig. S9, A and B**). On the other hand, we found a decrease in TBR2 labeling in MHS cerebral organoids, consistent with fewer intermediate progenitor cells compared to Ctrl and suggestive of a delay in development at this stage (**fig. S9, A and C**). There was also evidence for disruption of cortical layer formation in the MHS compared to Ctrl organoids, similar to that reported for conditional knockout of *MEF2C* in mouse brain (*12-15*). Along these lines, CTIP2, a cerebrocortical layer V-VI marker, and TBR1, a cortical layer VI marker, were markedly decreased in expression in MHS cerebral organoids compared to Ctrl (**fig. S9, A, D and E**).

In MEA recordings, we found a dramatic increase in electrical activity in MHS cerebral organoids compared to Ctrl (**Fig. 6, A, B and D**). Similar to our 2D cultures described above, MHS organoids showed increased frequency of network bursts and firing synchrony compared to Ctrl (**Fig. 6, E and F**). Importantly, NitroSynapsin normalized the excessive mean firing rate, network burst frequency, and firing synchrony of the MHS cerebral organoids virtually to Ctrl levels, while having minimal effect on Ctrl organoids (**Fig. 6, C to F**).

**Fig. 6.**
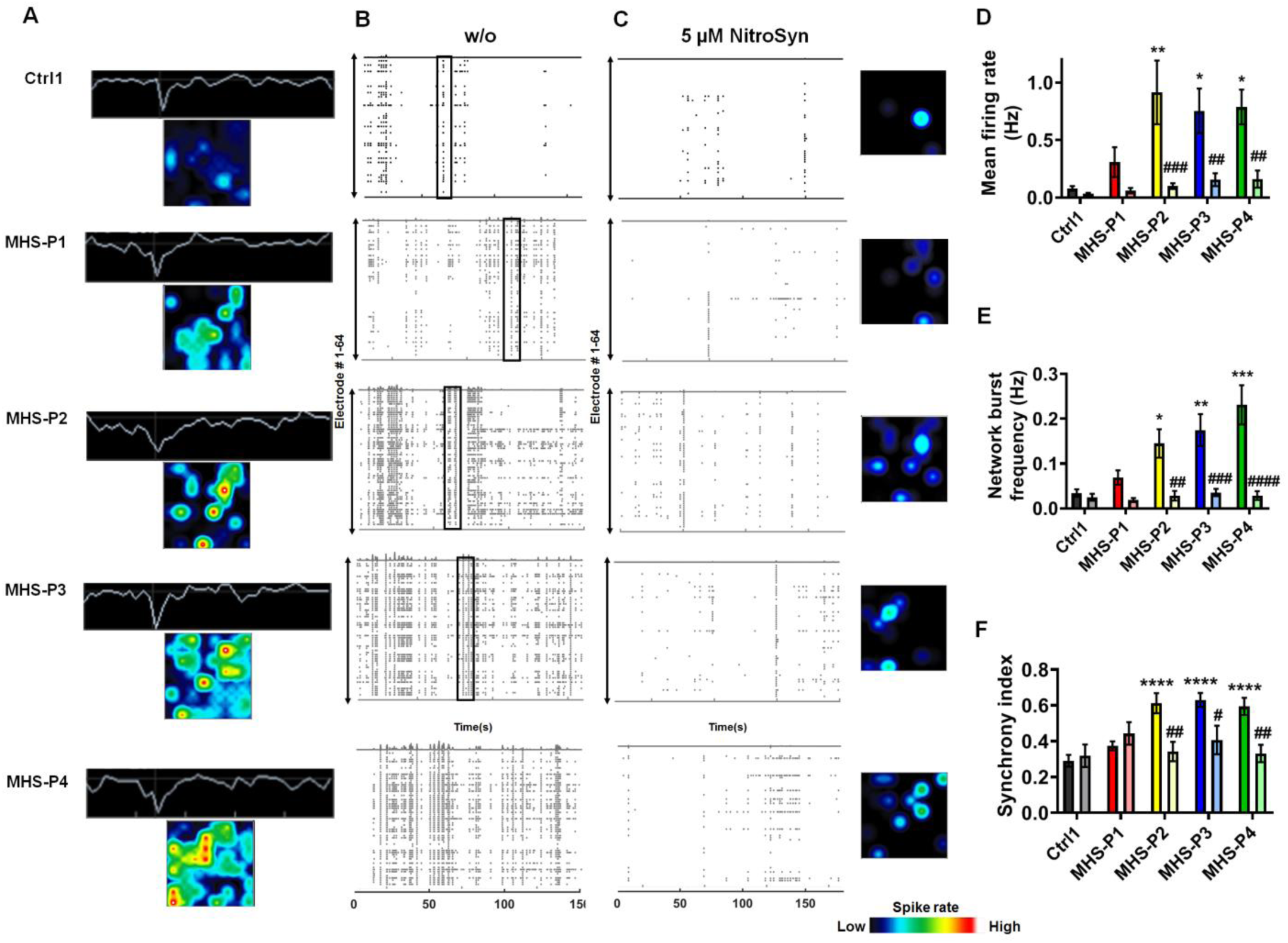
NitroSynapsin abrogates hypersynchronous burst activity in MHS cerebral organoids. **(A)** Representative heat maps and single traces from Ctrl1 and MHS hiPSC-derived cerebral organoids in individual MEA wells at 3-4 months of age. **(B)** Representative raster plots of MEA recordings in Ctrl and MHS cerebral organoids. Boxes outline examples of network bursts. **(C)** Representative raster plots and heat maps of MEA recordings in Ctrl and MHS cerebral organoids after treatment with NitroSynapsin (NitroSyn). **(D-F)** Quantification of MEA mean firing rate, network burst frequency, and synchrony index. Data are mean ± SEM. Sample size is listed above bars from 6-7 separate cerebral organoids recorded for each genotype. Ctrl includes data from Ctrl1. *^,#^*P* < 0.05, **^,##^*P* < 0.01, ***^,###^*P* < 0.001, ****^,####^*P* < 0.0001 by ANOVA for comparison with Ctrl value (*), or within a group (^#^) for without treatment (w/o) vs. NitroSynapsin treatment (see Methods).

## Discussion

As demonstrated in murine models, MEF2C plays a critical role during development to shape neuronal connectivity (*15, 31*). At early developmental stages, MEF2C is important for transcription of neuronally restricted genes and for the differentiation of hNPCs into neurons (*12*). In addition, MEF2C regulates formation and spread of dendrites, contributing to dendritic complexity and synapse formation, ensuring structural integrity of the neural network (*12, 15, 31, 65*). In contrast, at slightly later stages of development onward, MEF2C plays an important role in excitatory synapse pruning, promoting the formation and maintenance of the adult neural network (*50, 66*). Interestingly, heterozygosity for *MEF2C* in mice, mimicking human MHS patients, results in increased electrical excitability (*31*). Here, for the first time using patient-derived hiPSCs bearing MEF2C mutations, we investigated the effect of MEF2C on developing human neurons in the context of the MHS form of ASD/ID. We show that MEF2C haploinsufficiency yields cultures with more astrocytes and fewer neurons, at least in part due to aberrant miRNA expression. Furthermore, we found that while the total number of excitatory synapses was decreased on histological studies in MHS hiPSC-neurons compared to controls, there was an increase in VGLUT1 vesicular protein expression in excitatory presynaptic terminals and increased mEPSC frequency. These findings are also reflected in the increased excitability that we observed in network recordings from MHS hiPSC-neurons, in some cases resembling MHS patient seizures (**movie S2**). MHS hiPSC-neurons also displayed an increase in extrasynaptic activity, specifically, mediated by eNMDARs, consistent with overflow of neurotransmitter from hyperactive excitatory synapses (schematically summarized in **fig. S10**).

In mice, MEF2C has also been shown to regulate genes involved in inhibitory synapse formation (*51*), and is essential for the development of PV-expressing inhibitory neurons (*49*). In MHS hiPSC-neurons, we found decreased expression of the inhibitory vesicular protein VGAT accompanied by a decrease in mIPSC frequency, revealing dysfunctional inhibitory synapse activity. Along similar lines, systemic MEF2C haploinsufficiency in mice (with lower MEF2 activity as seen in human MHS patients) resulted in decreased mIPSC frequency (*31*). Seemingly in contrast to these findings, conditional knockout of *MEF2C* in embryonic cortical excitatory neurons (using *Emx1*^*Cre/+*^ knock-in mice crossed with *MEF2C*^*fl/fl*^ mice) led to a moderate decrease in excitatory transmission and a large increase in inhibitory transmission (*17*). However, those findings were the result of excitatory neuron-specific deletion of *MEF2C*, while inhibitory neurons were spared, since cortical excitatory neurons and glia but not GABAergic neurons are produced in the Emx1-expressing lineage (*67*). Notably, a similar decrease in excitatory synaptic activity was observed with conditional brain knockout of *MEF2C* in *Nestin-Cre*^*+*^*/MEF2C*^fl/Δ2^ conditional null mice (where Δ2 represents exon 2-deleted allele) (*12*), arguing that this phenotype is due to complete loss of MEF2C activity at a very early stage of development.

As found in the present study in MHS hiPSC-derived human neurons, where approximately half of MEF2C activity is lost, the increase in excitatory synaptic and extrasynaptic activity as well as the decrease in inhibitory synaptic transmission disrupts the excitatory-inhibitory (E/I) ratio. This contributes to hypersynchronous burst activity in neural networks. Several studies have associated E/I imbalance with ASD (*31, 68-70*). Critically, as in *MEF2C* heterozygous mice (*31, 68*), we found that the eNMDAR-preferring antagonist, NitroSynapsin, abrogated the aberrant network hyperactivity in MHS hiPSC-derived neuronal cultures and cerebral organoids, suggesting that this drug merits further study for its therapeutic potential. Moreover, since MEF2C has been shown to regulate expression of many hub genes known to be associated with ASD/ID (*21, 22, 43*), a similar approach may prove effective in these other forms of ASD/ID.

## Acknowledgments

The authors wish to thank Scott R. McKercher and other members of the Lipton laboratory for helpful discussions and insight. Fig. 2D and fig. S10 were created with BioRender.com.

## Author contributions

Conceptualization: D.T, S.G., R.A., S.A.L.

Investigation: D.T, S.G, J.P. M.T, T.G., S.M.N., M.T, M.L., N.D., C.B., K.L., A.S., S.F.C.

Formal analysis: D.T., S.G., A.C., Y.C., R.K., R.A., S.A.L.

Resources: P.S., I.B., P.K.

Writing – original draft preparation: D.T, S.G., S.A.L.

Writing – review and editing: D.T, S.G., S.A.L

Supervision: N.S., M.G.R., N.N., D.H.G., R.A., S.A.L.

Funding acquisition: S.A.L.

## Materials and Methods

### hiPSC reprogramming and maintenance

The use of human cells was approved by the institutional review boards of the Scintillon Institute and The Scripps Research Institute (TSRI; IRB-19-7428). hiPSCs were generated from patient fibroblasts, by using an integration-free reprogramming method (*71*), which uses three episomal vectors that collectively encode OCT3/4, SOX2, KLF4, L-MYC, LYN28, and p53-shRNA. hiPSCs were characterized for pluripotency, karyotypic integrity and generation of all three germ layers. hiPSCs were routinely cultured and maintained in our laboratory using a protocol described previously (*33*) with some modifications. Briefly, hiPSCs were plated on Matrigel coated plates (Corning, #354248) and cultured using mTeSR1 (STEMCELL Technologies, #05850) changed daily. The colonies were manually passaged weekly, using StemPro™ EZPassage™ Disposable Stem Cell Passaging Tool (Thermo-Fisher Scientific, #23181010).

### Generation of MEF2C deletion by CRISPR/Cas9

Ctrl1 hiPSCs (Hs27, ATCC #CRL-1634; see **table S1**) were used to produce the MEF2C deletion isogenic line at the Yale Stem Cell Center, which used CRIPSR/Cas9 to generate an 11bp deletion in MEF2C. Single clones were picked and further sequenced by Sanger sequencing. An isogenic line with an 11bp deletion was further characterized by whole-exome gene sequencing. No off-target effects were detected in coding regions. The isogenic line also maintained its pluripotency.

### Neuronal differentiation

Differentiation of hiPSCs was performed using standard protocols for generating cerebrocortical neurons (*33, 34*). Briefly, feeder-free hiPSCs cultured on Matrigel were induced to differentiate by exposure for 2 week to a cocktail of small molecules: 2 μM each of A83-01, dorsomorphin, and PNU74654 in DMEM/F12 medium supplemented with 20% Knock Out Serum Replacement (Invitrogen). Cells were then scraped manually to form floating neurospheres and maintained for 2 weeks in DMEM/F12 medium supplemented with N2 and B27 (Invitrogen) and 20 ng ml^−1^ of basic FGF (R&D Systems). Subsequently neurospheres were seeded on polyornithine/laminin-coated dishes to form rosettes that were manually picked, expanded and frozen down as human neural progenitor cells (hNPCs). For terminal differentiation into neurons, hNPCs were plated onto polyornithine/laminin-coated glass coverslips or plates at a density of 1.5×10^6^ cells/cm^2^ in DMEM/F12 medium supplemented with N2 and B27. For the first 2 days, 0.1 µM of compound E (γ secretase inhibitor, CAS 209986-17-4, Calbiochem) was added. Starting on the third day, the medium was supplemented with GDNF (20 ng ml^−1^) and BDNF (20 ng ml^−1^).

For electrophysiological analysis and RNA-seq experiments, hNPCs were plated at a 1:1 ratio with neonatal mouse astrocytes onto polyornithine/laminin-coated glass coverslips in DMEM/F12 medium supplemented with B27, N2, GDNF (20 ng ml^−1^), and BDNF (20 ng ml^−1^) (Peprotech), and 0.5% FBS (Invitrogen). Prior to electrophysiology experiments, cells at week 3 of terminal differentiation were switched to BrainPhys medium (STEMCELL Technologies), and experiments were conducted after a total of 5–6 weeks of differentiation. Our protocol produced 8–15% inhibitory neurons, as monitored by immunocytochemistry with anti-GABA antibody (*33*).

#### hiPSC-derived cerebral organoid cultures

Cerebral organoids were generated as previously described (*61*). In brief, spheroids were generated from hiPSCs using SMAD inhibitors, dorsomorphin and SB-4321542 for 5 days. From day 6, spheroids were transferred to Neurobasal medium supplemented with EGF and FGF-2 for 19 days. From day 25, the medium was supplemented with BDNF and NT3 until day 43, and from then on, the organoids were maintained in Neurobasal medium only. Cerebral organoids were maintained on an orbital shaker until used in experiments as described below. Histology was performed after 2 to 3 months of differentiation. At 3-4 months of age, the cerebral organoids were used for multielectrode array (MEA) analysis.

### Primary astrocytes

Primary astrocytes cultures were prepared as previously described (*72*). Mice used for astrocyte cultures were housed and maintained in the animal facility at the Scintillon Institute or The Scripps Research Institute, and all experiments complied with protocols approved by the Institute Animal Care Committee (IACUC protocol, 17-0022-2). The brains from 1-to 2-day-old C57BL/6 mouse pups were rapidly harvested, and the meninges were removed. Brains were dissociated using mechanical means (pipettes and scissors) and enzymatic dissociation (0.25% <u>trypsin</u> for 30 min at 37 °C). The cell suspension was then filtered through a 70-µm cell strainer and cultured in DMEM:F12 medium supplemented with 10% <u>fetal bovine serum</u>, 100 U/ml <u>penicillin</u>, and 0.1 mg/ml <u>streptomycin</u>. Two weeks after initial isolation, cells were dissociated with Accutase (STEMCELL Technologies, #07920), and the suspension was seeded onto glass coverslips coated with 20 µg/cm^2^ poly-L-ornithine (Sigma-Aldrich, #P4638), 2 µg/cm^2^ laminin (Trevigen, #3400-010-01), and 2 µg/cm^2^ fibronectin (Trevigen, #3420-001-01) at a density of 25,000 cells/cm^2^.

### Immunoblots

Cells were lysed with cell lysis buffer (Cell Signaling Technologies, #9803) supplemented with protease inhibitor cocktail (Roche, #04693159001) and 10 µM SDS. Protein concentrations were determined using Pierce™ BCA Protein Assay Kit (Thermo-Fisher Scientific, #23225). Polyacrylamide gels (Novex, 8-12%) were used for protein separation prior to transfer onto nitrocellulose membrane, blockade with blocking buffer (Li-Cor) for 1h, washing with TBST, and reaction overnight at 4 °C with mouse anti-MAP2 (1:2,000; Sigma-Aldrich, #M4403, RRID:AB_477193) and then mouse anti-GAPDH (1:10,000; Sigma-Aldrich, #CB1001 EMD MILLIPORE, RRID:AB_2107426). Membranes were then reacted with goat anti-mouse infrared (IR) dye-conjugated secondary antibody (Li-Cor antibody, #926-32210, RRID AB_621842) for 1 h at room temperature (RT). The membrane was scanned using an Odyssey® scanner, quantified using Fiji software, and analyzed with Prism 7.04 software (GraphPad).

### Transfection with miRNA inhibitor

Ctrl1 hNPCs were transfected with miRNA inhibitors using HiPerFect Transfection reagent (Qiagen, # 301705) according to the manufacturer’s instructions. The following inhibitors were used (purchased from Qiagen): hsa-miR-663b miRCURY LNA miRNA Inhibitor (YI04100013); hsa-miR-4273 miRCURY LNA miRNA Inhibitor (YI04105965); negative control A miRCURY LNA miRNA Inhibitor Control (YI00199006). After 48 h, the medium was replaced with fresh medium, and cells were maintained for an additional 2 weeks before being fixed and stained.

### Quantitative (q)RT-PCR

For total RNA, RNA was extracted using a Quick-RNA™ MiniPrep kit (Zymo Research, #R1055), and each sample was reverse transcribed using a QuantiTect Reverse transcription kit (Qiagen, #205313). qRT-PCR reactions were performed with LightCycler 480 SYBR Green I MasterMix (Roche) in a LightCycler480II instrument (Roche). The primer sequence was designed using Primer Bank (*73*). NRXN3: F-AGTGGTGGGCTTATCCTCTAC; R-CCCTGTTCTATGTGAAGCTGGA. S100β: F-TGGCCCTCATCGACGTTTTC; R-ATGTTC AAAGAACTCGTGGCA. The following primers were purchased from Qiagen: SOX2, NANOG, Lin28, β3-tubulin. Data were normalized to 18s rRNA expression, and analyzed with Prism 7.04 software (GraphPad).

miRNA was extracted using a mirVana™ miRNA Isolation Kit (Thermo Fisher, #AM1560), and each sample was reverse transcribed using miScript II RT Kit (Qiagen, #218161). qRT-PCR reactions were performed with miScript SYBR Green PCR Kit (Qiagen, #218075) in a LightCycler480II instrument (Roche). The following primers were purchased from Qiagen: miR-663b (MS00037884), miR-663 (MS00037247), miR-4273 (MS00021280). Data were normalized to RNU6-2 expression, and analyzed with Prism 7.04 software (GraphPad).

### Histological imaging and immunocytochemistry

For 2D cultures, hiPSC colonies, hNPCs, or neurons at 2 time-points of differentiation (1 and 3 months) were fixed with 4% PFA for 15 min and washed 3 times with PBS. Phase-contrast images of 2-month-old 3D organoids were acquired on an EVOS cell imaging system (Thermo-Fisher Scientific) for size measurements. Two to three-month-old cerebral organoids were fixed in 4% PFA at 4 °C overnight followed by serial incubation in 15% and 30% sucrose in PBS overnight. Fixed organoids were embedded in tissue freezing medium (TFM, General Data, Cincinnati, OH), and were flash frozen with isopentane and liquid nitrogen. Frozen organoids were sectioned in a cryostat using optimal cutting temperature (OCT) compound and then sectioned at 15-µm thickness. Cells or sections were blocked with 3% BSA and 0.3% Triton X-100 in PBS for 30 min. For synaptic staining, cultures were blocked with 3% BSA and 0.1% Saponin (wt/vol) in PBS. Cells/sections were incubated with primary antibody in blocking solution overnight at 4 °C and then washed with PBS. The appropriate Alexa Fluor (488, 555, 647) conjugated secondary antibodies were used at 1:1,000, plus Hoechst 33342, Trihydrochloride, Trihydrate dye (1:1,000, Thermo-Fisher Scientific, #H3570) to visualize nuclei for 1 h at RT.

Primary antibodies and dilutions were as follows: Rabbit anti-NANOG (1:500, Cell signaling technologies, #4903s, RRID:AB_10559205); mouse anti-TRA1-60 (1:500, Cell Signaling Technologies, #4746s, RRID:AB_2119059); rabbit anti-SOX2 (1:500, Cell Signaling Technologies, #3579s, RRID:AB_2195767); mouse anti-Nestin (1:250, Abcam, #ab22035, RRID:AB_446723); mouse anti-S100β (1:1,000, Abcam, #ab11178, RRID:AB_297817); rabbit anti-GFAP (1:500, Agilent, #Z0334, RRID:AB_10013382); mouse anti-MAP2 (1:500, Sigma-Aldrich, #M4403, RRID:AB_477193); chicken anti-β3-tubulin (1:500, Abcam, #ab41489, RRID:AB_727049); guinea pig anti-VGLUT1 (1:250, Synaptic Systems, #135304, RRID:AB_887878); mouse anti-VGAT (1:250, Synaptic Systems, #131011, RRID:AB_887872); rabbit anti-synapsin I (1:500, Millipore, AB1543p, RRID:AB_90757); mouse anti-PSD95 (1:500, Invitrogen, MA1-045, RRID:AB_325399); rat anti-CTIP2 (1:150; Abcam, #ab18465, RRID:AB_2064130); rabbit anti-TBR2 (1:300; Abcam, #ab23345, RRID:AB_778267); rabbit anti-TBR1 (1:250, Abcam, #ab31940, RRID:AB_2200219). Coverslips were mounted on slides with fluorescent mounting medium (DAKO) and visualized with a Zeiss Axiovert (100M) epifluorescence microscope or ImageXpress automated high-content confocal microscopy (Molecular Devices). For synaptic staining of PSD95 and synapsin I, punctae were visualized using a Nikon A1 confocal microscope. Imaging conditions were identical for each set of stains (e.g. exposure time and laser power). Quantification of GFAP, MAP2, nestin, TBR2, TBR1, CTIP2, synapsin I, and PSD-95 were performed with Fiji software. The number of synapses was calculated as co-localized punctae of synapsin I/PSD-95 staining per neurite length identified with β3-tubulin. Quantification of S100β, β3-tubulin, VGLUT1, and VGAT were performed using MetaXpress (Molecular Devices). Quantification of immunostained cells was performed in a masked fashion, normalized to cell number (by Hoechst staining for cell nuclei), and compared with the same masking/thresholding settings.

### Luciferase gene reporter assays

Luciferase reporter assay was performed as previously described with some modifications (*39*). Briefly, hNPCs were transfected with MEF2 luciferase reporter (*74*) along with *Renilla* luciferase control vector using a Human Stem Cell Nucleofector Kit according to the manufacturer’s instructions (Lonza, VPH-5012). Cells were plated in a 96-well plate and terminally differentiated into neurons. Cells were harvested at various time points of differentiation over a period of 11 days and then analyzed using a Dual-Glo luciferase assay kit (Promega) following the manufacturer’s instructions. Firefly luciferase activity was normalized to *Renilla* luciferase activity.

### RNA-seq data acquisition, mapping, and analysis

After 5 weeks in culture, hiPSC-derived neurons grown on mouse neonatal astrocytes were processed for analysis. The following patient samples were used for the RNA sequencing: control samples Ctrl1, Ctrl2, and Ctrl4, and patient mutations MHS-P1, MHS-P2, MHS-P3, and MHS-P4 (see **table S1**). RNA was extracted using a Quick-RNA™ MiniPrep kit (Zymo Research, #R1055). RNA was quality checked before sequencing by RNA screenTape (Agilent tape station) and nanodrop. All RNA samples had an RNA integrity number (RIN) >9. RNA libraries were prepared via Ribo depletion. Sequencing was performed on the Illumina HiSeq2500 platform in a 1×75 bp single end (SR) configuration in high output mode (V4 chemistry) with 40 million reads per sample. Reads were aligned to the latest human hg38 reference genome using the STAR spliced read aligner (*75*). No read trimming or filtering was done with this data set because the quality distribution and variance appeared normal. Total counts of read-fragments aligned to known gene regions within the human hg38 refSeq reference annotation were used as the basis for quantification of gene expression. Fragment counts were derived using HTS-seq program using hg38 Ensembl transcripts as model. Various QC analyses were conducted to assess the quality of the data and to identify potential outliers. Differentially expressed genes (DEGs) were identified using three bioconductor packages, EdgeR, Limma+Voom and Limma, which were then considered and ranked based on adjusted *P*-values (FDR) of ≤0.1. Gene set enrichment analysis (GSEA) was performed using the top DEGs (determined by EdgeR). DEGs were selected based on several false discovery rate (FDR) Benjamini-Hochberg adjusted *P*-values and simple *P*-values. A total of 1029 genes were used for the enrichment analysis. Of the neuronal-relevant GO terms, the most differentially expressed genes were further analyzed by qPCR to validate the results.

### ChIP-seq and qRT-PCR

ChIP assays were performed using the *ChIP-IT Express Kit* (Active Motif) according to the manufacturer’s instructions. **Briefly**, ∼2 × 10^7^ hNPCs were used per ChIP experiment. The hNPCs were first dissociated with Accutase and crosslinked with 1% paraformaldehyde, and then the nuclei were sonicated 10 times for 20 s at maximum settings using a Misonix Sonicator (Misonix, Farmingdale, NY). Sheared chromatin was immunoprecipitated with 4 μg MEF2 antibody (#sc-313, Santa Cruz Biotechnology, RRID:AB_631920) or control IgG (Active Motif) antibody. At this stage of development, hNPCs express primarily the MEF2C isoform of the MEF2 transcription factor family. ChIP DNA samples were subjected to qPCR assay. We determined the enrichment of specific DNA sequences after ChIP using an EXPRESS SYBR® GreenER™ detection kit (Invitrogen) on a Mx3000P real time PCR system (Stratagene). Levels of enrichment after ChIP were calculated using the comparative cycle threshold method (Invitrogen) after normalizing with IgG control, as we have previously described (*35, 36*). ChIP DNA samples were also subjected to preparation for ChIP-Seq Library according to Illumina’s Chip-Seq Sample prep kit. ChIP-seq samples were sequenced with an Illumina Genome Analyzer II system (San Diego, CA) according to the manufacturer’s instructions and aligned to the University of California, Santa Cruz (UCSC) *Homo sapiens* reference genome (hg18) using Bowtie. The aligned reads were processed with HOMER v2.6 software and visualized on the UCSC genome browser (*76*). Peaks were predicted and annotated with HOMER v2.6 software **(table S3)**.

### Electrophysiology and pharmacology experiments

Whole-cell recordings were performed with patch electrodes of 3 to 5 MΩ resistance. To analyze voltage-gated currents, pipettes were filled with an internal solution composed of (in mM): K-gluconate, 120; KCl, 5; MgCl_2_, 2; HEPES, 10; EGTA; 10; Mg-ATP, 4; pH 7.4 and mOsm 290. The external solution was composed of Ca^2+^ and Mg^2+^-free Hank’s Balanced Salt Solution (HBSS; GIBCO) to which were added: CaCl_2_, 2 mM; HEPES, 10 mM; and glycine, 20 mM. Recordings were performed using a Multiclamp 700B amplifier (Molecular Devices) at a data sampling frequency of 2 kHz with an analog-to-digital convertor, Digidata 1440A (Molecular Devices). Voltage-clamp and current-clamp protocols were applied using Clampex v10.6 (Molecular Devices). Preliminary analysis and offline filtering at 500 Hz were achieved using Clampfit v10.6 (Molecular Devices).

Agonist-induced currents (glutamate-and GABA-evoked) were recorded at a holding potential of −70 mV in the nominal absence of extracellular Mg^2+^ and in the presence of 20 μM glycine (Sigma) and tetrodotoxin (TTX, 1 μM, Hello Bio). Agonist was applied via a rapid gravity-flow, local superfusion system. The internal solution for these recordings was (in mM): CsCl, 135; MgCl_2_, 2; HEPES, 10; EGTA; 1; Mg-ATP, 4; pH 7.4 and mOsm 290.

Evoked postsynaptic currents were induced by brief (1 ms) unipolar current pulses (0.1–0.9 mA) from an external stimulator (A365, WPI) using a stimulating electrode (CBAEC75, FHC) positioned at a distance of 100–150 μm from the cell soma (*77*).

Spontaneous postsynaptic currents (sPSCs) were recorded at a holding potential of -70 mV and 0 mV in gap-free mode as described in previously (*33*). The internal solution contained (in mM): K-gluconate, 120; KCl, 5; MgCl_2_, 2; HEPES, 10; EGTA; 10; Mg-ATP, 4; pH 7.4 and mOsm 290. Miniature excitatory postsynaptic currents (mEPSCs) and miniature inhibitory postsynaptic currents (mIPSCs) were recorded at -70 mV and 0 mV, respectively, at 21 °C after equilibrium in TTX (1 µM) for at least 20 min. The internal solution used for recording mEPSCs and mIPSCs was comprised of (in mM): Cs-gluconate 130; CsCl 5; MgCl_2_, 2; HEPES, 10; EGTA; 1; Mg-ATP, 4; pH 7.4 and mOsm 290. Mini Analysis software (Synapstosoft, Fort Lee, NJ) was used to calculate the frequency and amplitude of spontaneous synaptic events.

Extrasynaptic currents were pharmacologically isolated and measured by whole-cell patch clamp recordings of hiPSC-derived cerebrocortical neurons based on previously published protocols (*56, 58*). Briefly, continuous recordings of individual neurons were performed in gap free mode at -70 mV in control conditions, followed by application of bicuculline (50 µM, a GABA_A_ receptor antagonist, Tocris) for 1 min to inhibit inhibitory synaptic inputs and activate synaptic glutamate receptors. Subsequently, the synaptic NMDARs thus activated were blocked by application of the open-channel irreversible antagonist MK-801 (10 µM, Tocris) in the continued presence of bicuculline for 5 min, followed by a 30 min wash with external solution. The residual current recorded after the wash was analyzed as extrasynaptic current. Thereafter, memantine or NitroSynapsin was applied to the neuron to study its effect on this current since these drugs have been shown to predominantly inhibit eNMDARs (*32, 53, 55*).

### Calcium imaging

For monitoring the relative change in intracellular Ca^2+^ concentration vs. time, Fluo-4 (Fluo 4 AM/Fluo 4 direct, Life Technologies) was used according to the manufacturer’s instructions and a previously published protocol (*54*). To monitor Fluo-4 fluorescence changes, randomly selected fields of comparable cell density were excited at 480 nm with an exposure time of 30 ms, imaged using a 40X oil objective at 33 frames per second, and analyzed with MetaMorph software (Molecular Devices). Changes in intracellular Ca^2+^ from baseline (F_0_) to peak fluorescence (ΔF) were expressed as fractional changes above baseline (ΔF/F_0_). Events with ΔF/F rise times of <200 ms were considered neuronal transients. Neurons were distinguished from astrocytes morphologically, as verified with cell-type specific antibody staining in previously published experiments (*33*). Calcium imaging movies using Fluo-4 were prepared at 500 frames per second using ImageJ (NIH).

eNMDAR-mediated Ca^2+^ changes were isolated using the previously published protocol with bicuculline and MK-801 delineated above (*56*). After washing out the MK-801 for 30 min, subsequent changes in intracellular Ca^2+^ levels represented eNMDAR activity, which could be blocked by NitroSynapsin (*32, 53*).

### Multielectrode array (MEA) recording

MEA recordings were obtained as previously described (*52*). Cells were plated on CytoView 12-well plates (Axion Biosystems) coated with 0.1% polyethyleneimine (PEI) and 10 μg/ml laminin. Based upon pilot experiments performed to determine when neuronal properties developed, we recorded from 2D cultures 5–6 weeks after neuronal differentiation from hNPCs, and from cerebral organoids at 3-4 months of development. Recordings were performed on a Maestro MEA (Axion Biosystems) using the “neural broadband analog mode” setting with a sampling frequency of 12.5 kHz in control conditions and after NitroSynapsin treatment at 37 °C and analyzed with Axion Biosystems Maestro Axis Software (version 2.4.2). For calculating firing frequency, an electrode displaying >5 spikes per min was considered active. A network burst was defined as a minimum of 25 electrodes displaying >10 spikes per electrode at <100 ms inter-spike interval. Network burst frequency was calculated as total number of network bursts recorded per second during the analysis window expressed in Hz. Synchronous firing was determined by analyzing a 20 ms synchrony window, which is the time window around zero used for computing the area under the cross-correlation curve. Analysis of well-wide, pooled interelectrode cross-correlation gives the synchrony index, defined as a unitless measure of synchrony between 0 and 1 such that values closer to 1 indicate higher synchrony.

### Data analysis and statistics

For each experiment, we used at least 3 independent sets of cultures from separate differentiations or 3 independent groups of cerebral organoids from separate differentiations. Sample size was determined from prior power analyses based on prior data obtained in our laboratory. All data were acquired by an investigator blinded to the sample groups. No data was excluded from analysis. Data are presented as mean ± SEM. Statistical analyses were performed using GraphPad Prism software. Statistical significance was determined by a two-tailed unpaired Student’s t test for single comparisons, by ANOVA followed by a post hoc Dunnett’s test for multiple comparisons, or by a Sidak’s test corrected for multiple comparisons between selected pairs. For non-parametric data such as percentage change in calcium event frequency, we used a Kruskal-Wallis test followed by a post hoc Dunn’s test for multiple comparisons. For analysis of cumulative distributions, we used the Kolmogorov-Smirnov test performed with Mini Analysis software (Synapstosoft, Fort Lee, NJ). Data with *P* values < 0.05 were considered to be statistically significant. In general, the variance was similar between groups being compared. No samples or data were excluded from the analysis.

## Supplementary Figures

**Fig. S1.**
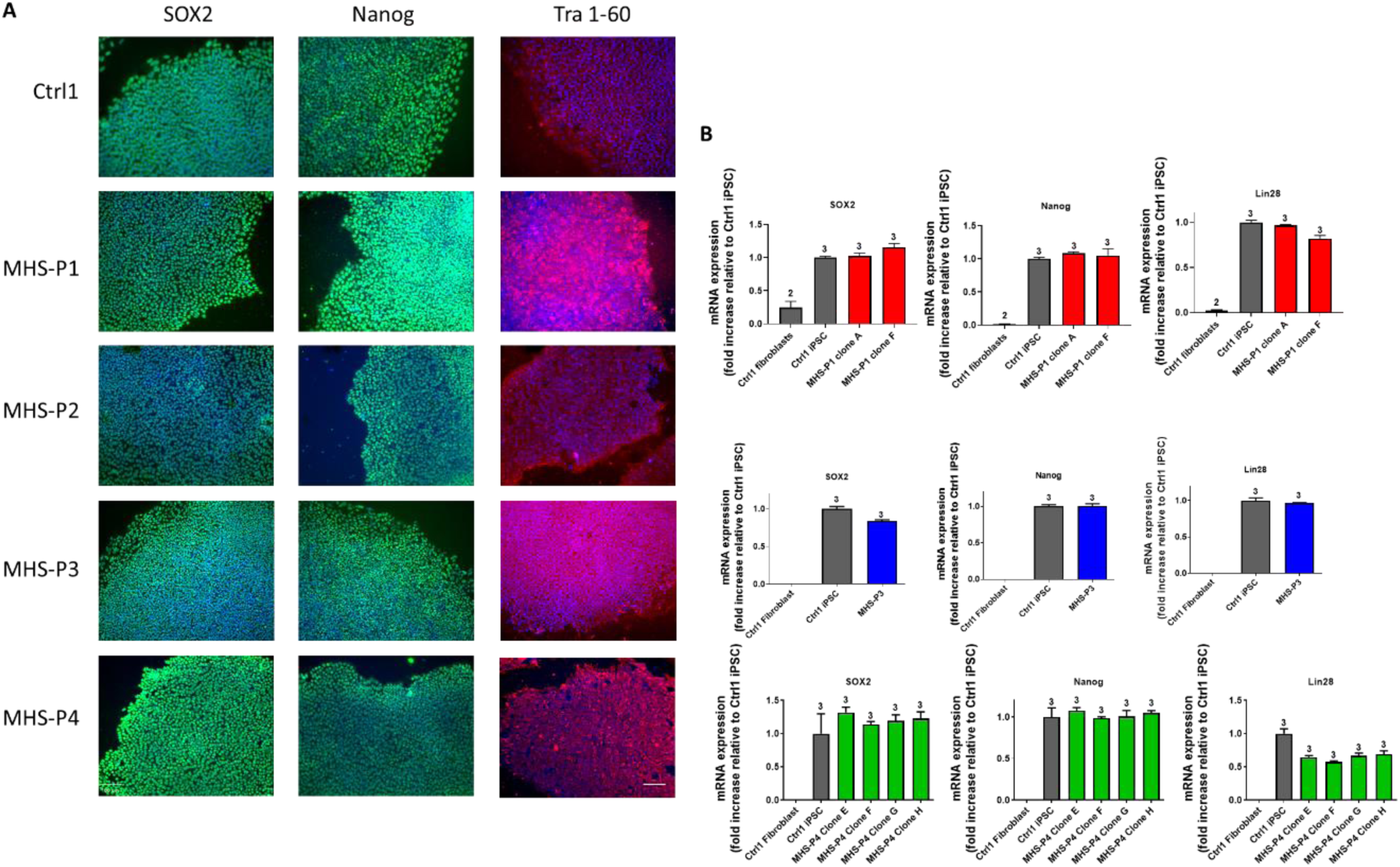
Generation of hiPSCs from patient fibroblasts. **(A)** Immunocytochemistry showing expression of pluripotency markers SOX2, Nanog and Tra1-60 in all hiPSC lines. Scale bar, 100 µm. **(B)** qPCR analysis of pluripotency markers SOX2, Nanog and Lin28 for several clones generated for each MHS patient hiPSC vs. Ctrl fibroblasts. Data are mean + SEM. Sample size listed above bars (number of replicates).

**Fig. S2.**
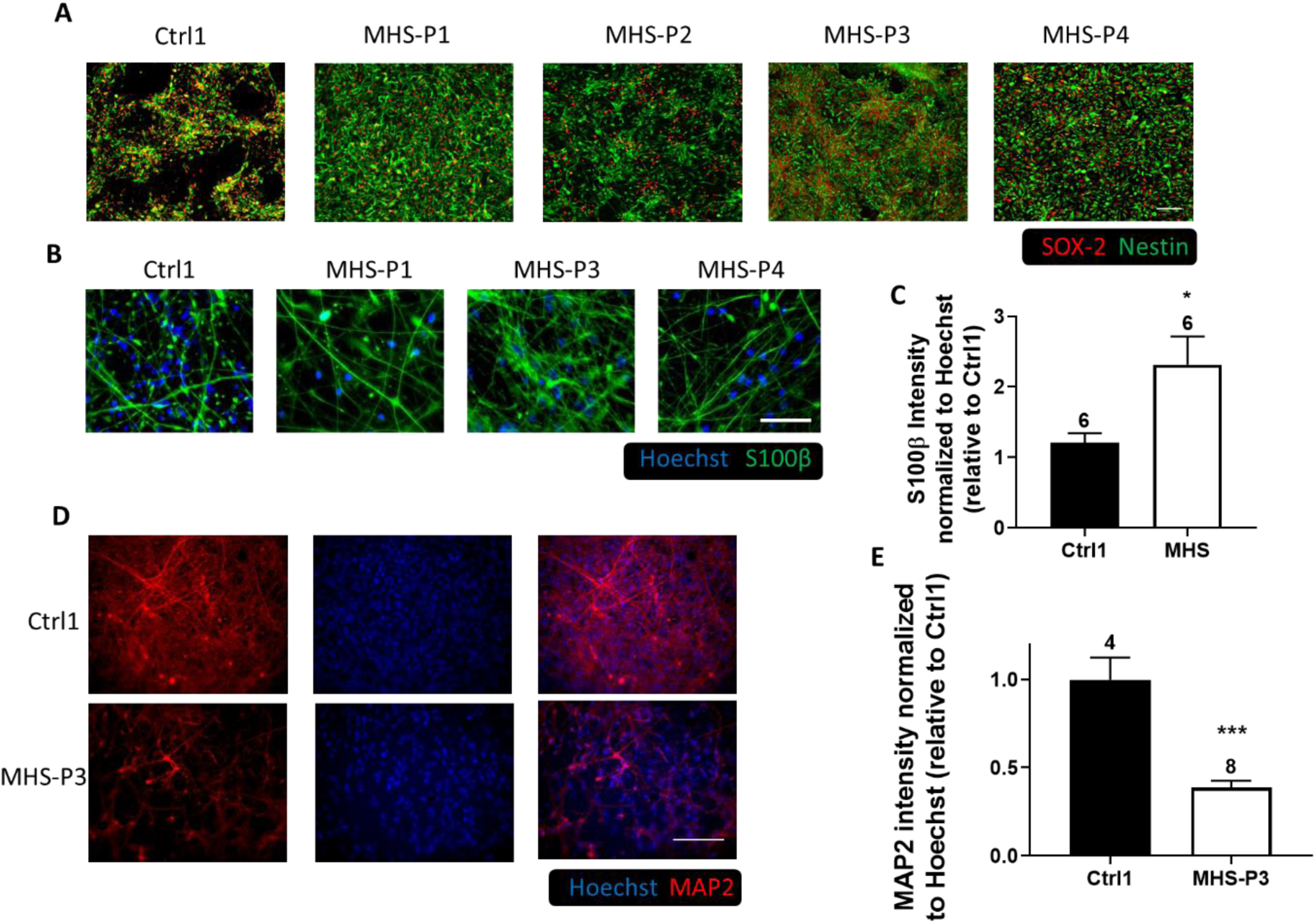
MHS patient hNPC characterization. **(A)** Immunocytochemistry of SOX2 and Nestin markers in Ctrl and MHS patient hiPSC-derived hNPCs. Scale bars, 100 µm. **(B)** Immunocytochemistry of S100β and Hoechst in neurons differentiated from Ctrl and MHS hiPSCs after 1 month. Scale bars, 50 µm. **(C)** Quantification of S100β expression. **(D)** Immunocytochemistry of MAP2 in Ctrl1 and MHS-P3 patient hiPSC-derived neurons at 3 months. Scale bars, 100 µm. **(E)** Quantification of MAP2 expression in 3-month-old neurons from Ctrl and MHS. Data are mean + SEM. Sample size listed above bars (number of replicates in 3 independent experiments). \**P* < 0.05, \*\*\**P* < 0.001 by Student’s t test.

**Fig. S3.**
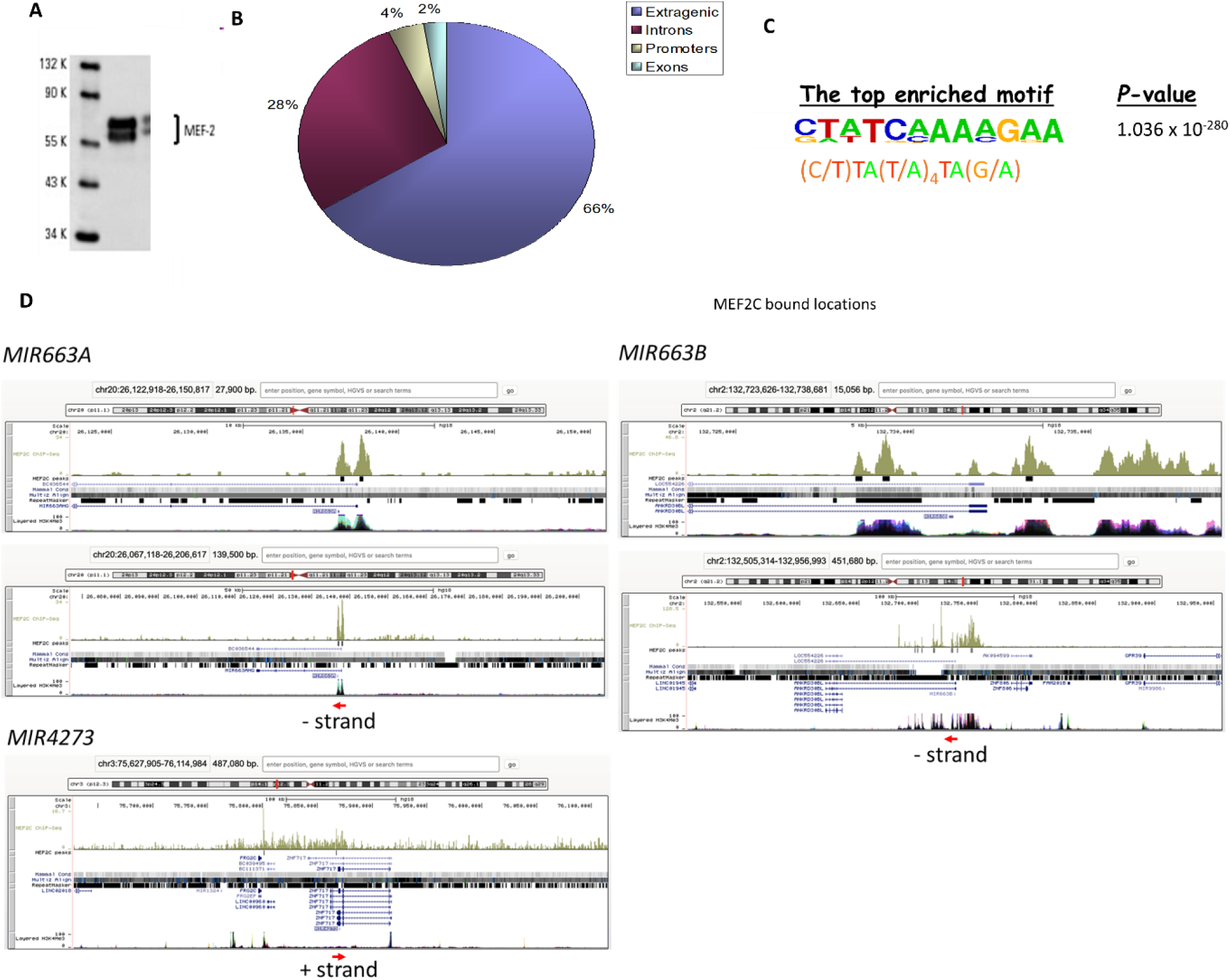
ChIP-seq for MEF2 targets. **(A)** Validation of MEF2 antibody, showing specific binding to MEF2 (including MEF2A, C and D). At this time of development, however, primarily MEF2C is expressed. **(B)** Pie chart showing the distribution of MEF2 binding sites on the genome. Blue indicates extragenic sequences that bind MEF2. (**C**) Top-enriched motif for MEF2 binding. **(D)** ChIP-seq data sets for miR663a, miR663b and miR4273 in the vicinity of the *MEF2C* gene locus.

**Fig. S4.**
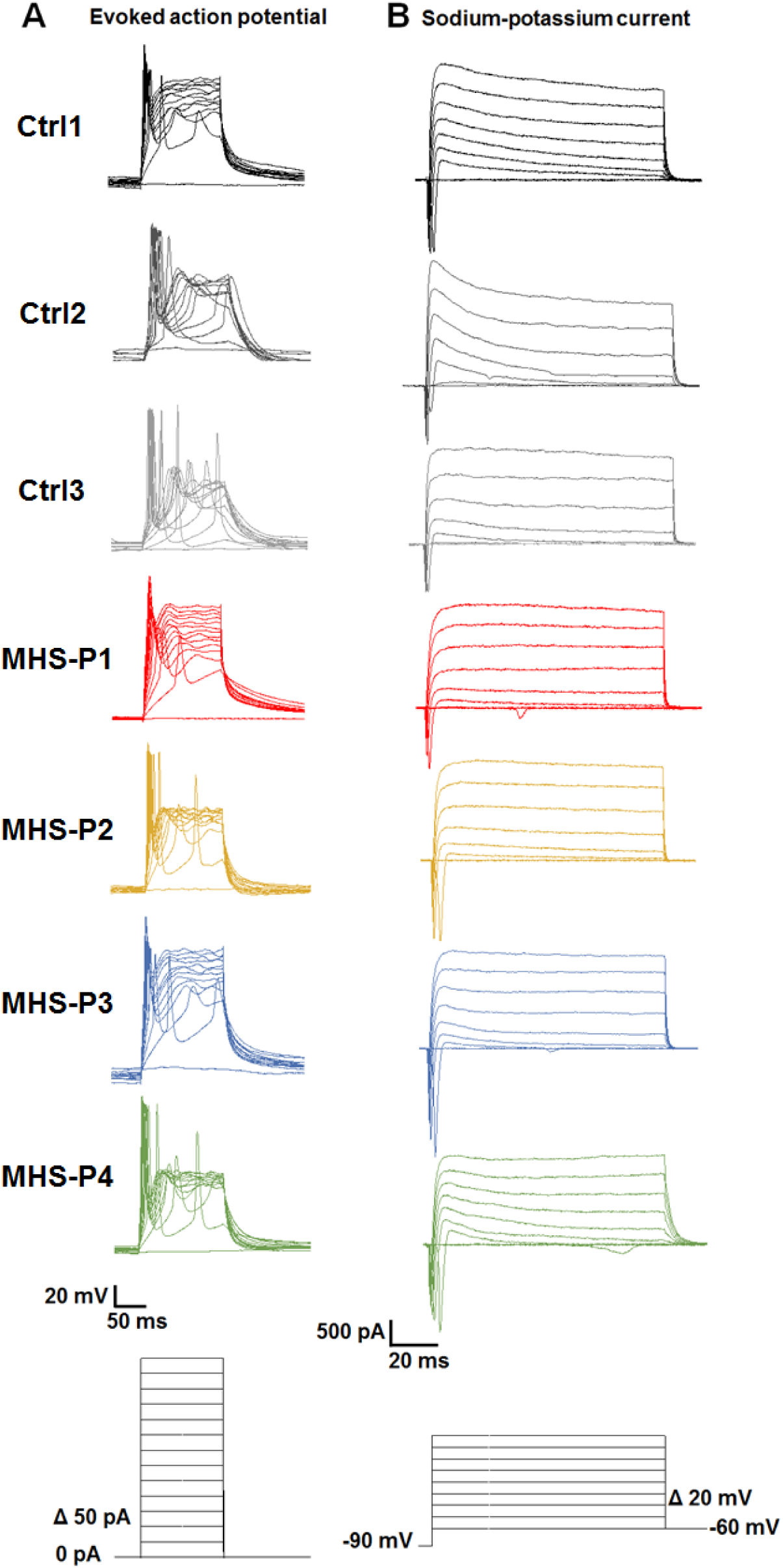
Both Ctrl and MHS hiPSC-derived cerebrocortical neurons fire action potentials (APs) in current clamp, and manifest sodium and potassium currents under voltage-clamp. **(A)** Patch-clamp recording of evoked APs in hiPSC-derived neurons from a holding potential (V_h_) of -60 mV in current-clamp mode. Current-injection protocol used to evoke APs illustrated below traces. **(B)** Representative sodium and potassium currents in voltage-clamp mode elicited by voltage steps from V_h_ = -60 mV after a prepulse to -90 mV for 300 ms. Voltage protocol illustrated below traces.

**Fig. S5.**
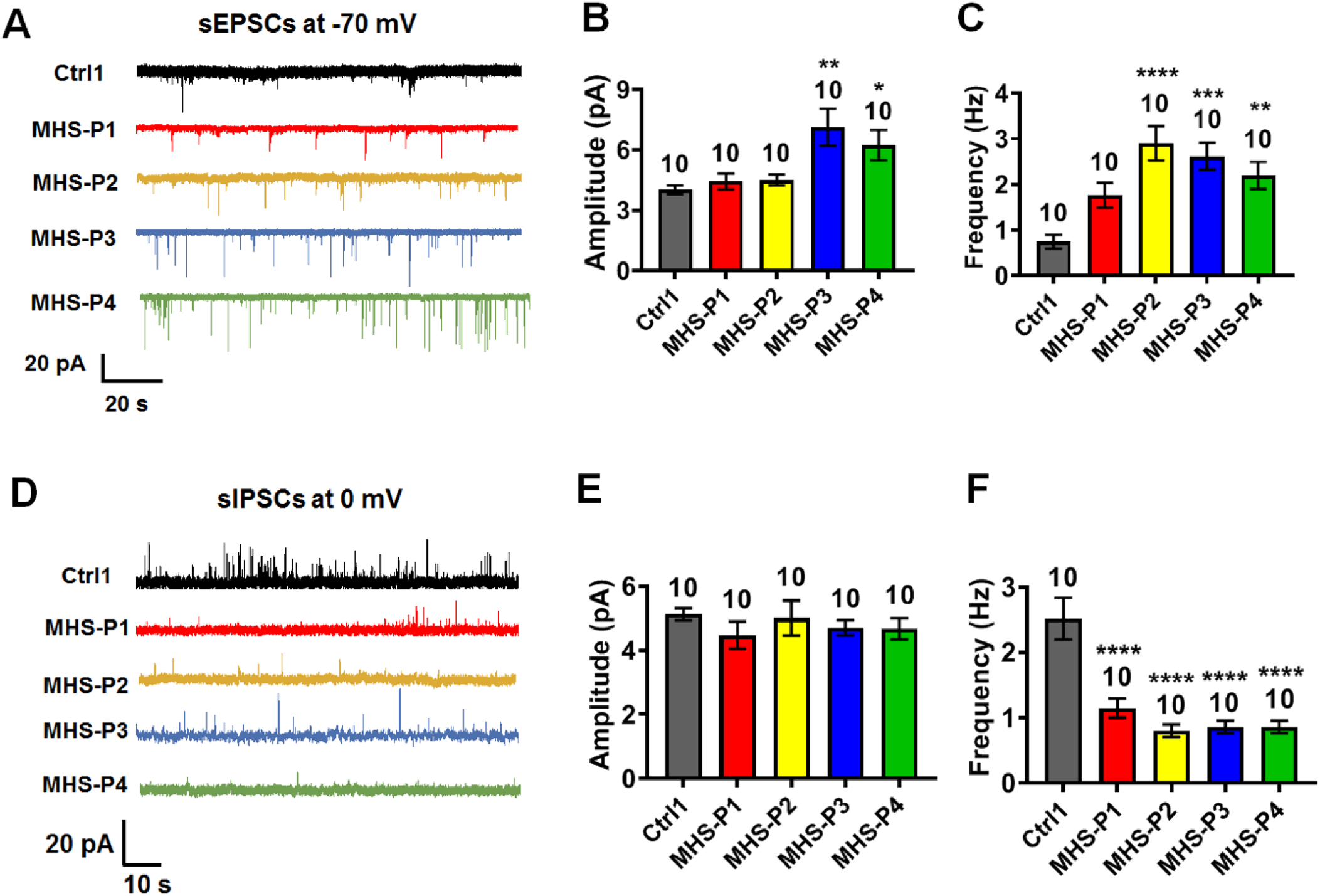
MHS hiPSC-derived cerebrocortical neurons exhibit increased excitatory synaptic activity and decreased inhibitory synaptic activity. **(A)** Representative spontaneous (s)EPSCs recorded at -70 mV from Ctrl and MHS hiPSC-neurons in culture for 5 weeks. **(B** and **C)** Quantification of sEPSC mean amplitude and frequency. **(D)** Representative sIPSCs recorded at 0 mV from Ctrl and MHS hiPSC-neurons in culture for 5 weeks. **(E** and **F)** Quantification of sIPSC mean amplitude and frequency. Data are mean ± SEM. Sample size (*n*) listed above bars from 3-5 experiments. \**P* < 0.05, \*\**P* < 0.01, \*\*\**P*<0.001, \*\*\*\**P* < 0.0001 by ANOVA with Dunnett’s post hoc test for multiple comparisons.

**Fig. S6.**
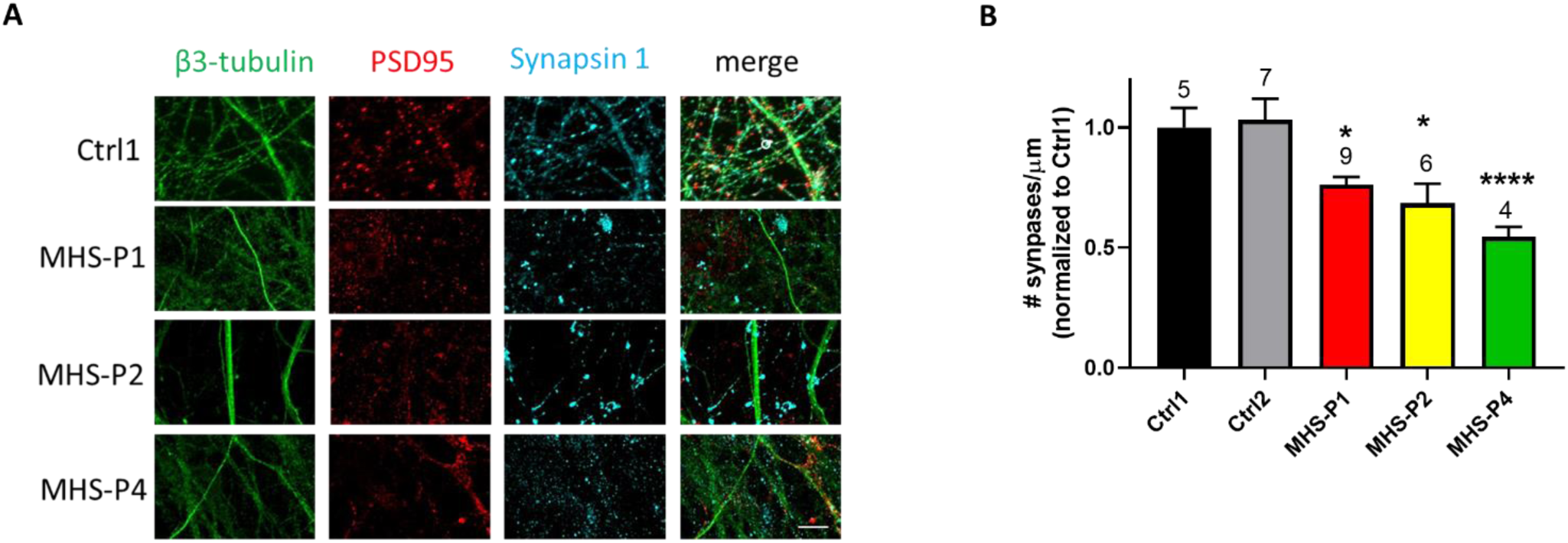
MHS hiPSC-derived cerebrocortical neurons exhibit fewer synapses. **(A)** Representative images of β3-tubulin, PSD-95, synapsin I, and merged image for Ctrl and MHS patients. Scale bar, 10 µm. **(B)** Quantification of number of synapses (coincident synapsin 1/PSD-95 punctae staining) per neurite length for Ctrl and MHS patients. Data are mean + SEM. Number of imaged fields (*n*) listed above bars from 3 separate experiments. \**P* < 0.05, \*\*\*\**P* < 0.0001 by ANOVA with Dunnett’s post hoc test for multiple comparisons.

**Fig. S7.**
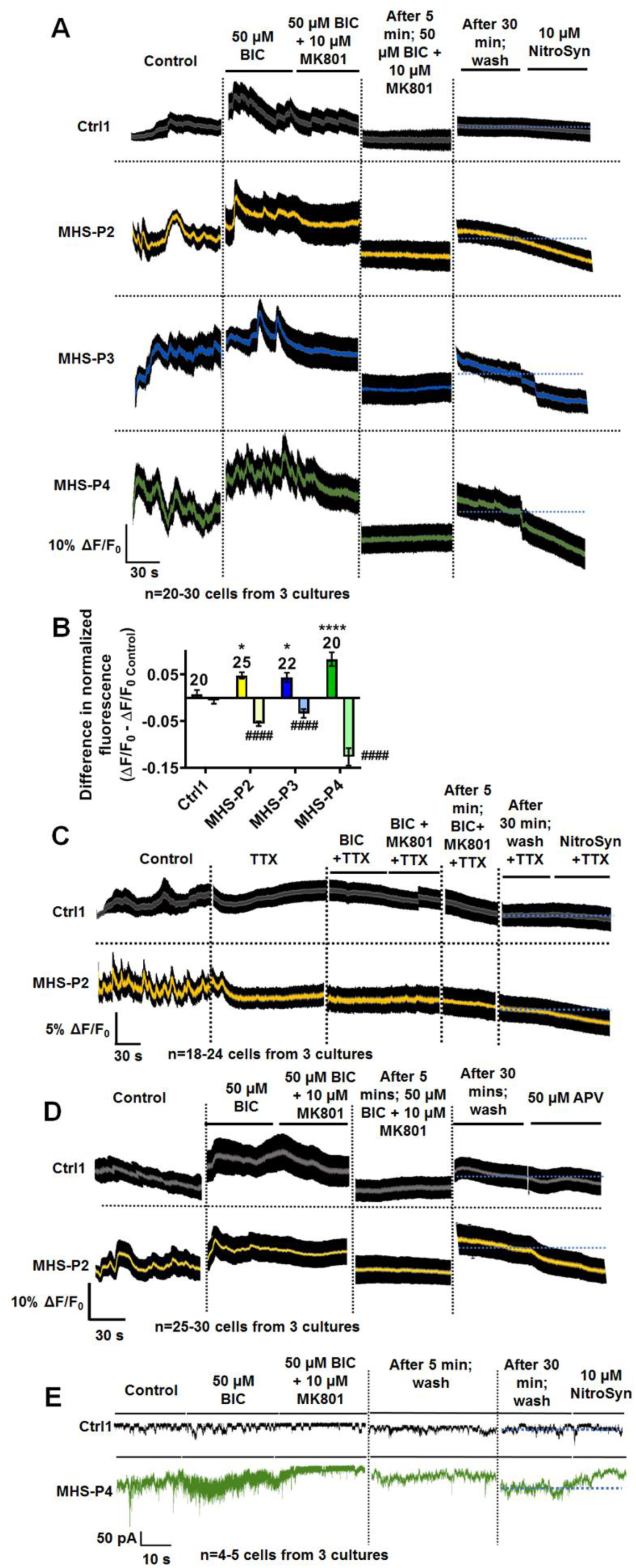
NitroSynapsin inhibits aberrant basal eNMDAR-responses in MHS hiPSC-derived cerebrocortical neurons. **(A)**. Mean ± SEM envelope of calcium responses and their inhibition by NitroSynapsin (NitroSyn) monitored with Fluo-4 in Ctrl and MHS neurons after pharmacological isolation of eNMDARs. The protocol for isolating eNMDAR responses consisted of activation and subsequent blockade of synaptic NMDAR-mediated currents by bicuculline (BIC) and MK-801, respectively. Blue-dotted line added to show Ca^2+^ response before and after NitroSynapsin. **(B)** Quantification of basal eNMDAR-mediated Ca^2+^ responses and effect of NitroSynapsin. Reponses before treatment (bright bars with positive values, signifying the amplitude of eNMDAR-mediated response) and after treatment (NitroSynapsin; pale bars with negative values, representing inhibition of the eNMDAR-mediated response). **(C)** Mean ± SEM envelope of calcium traces from Ctrl and MHS hiPSC-neurons after pharmacological isolation of eNMDAR-mediated responses in the presence of TTX. **(D)** Mean ± SEM envelope of calcium traces showing inhibition with APV in Ctrl and MHS neurons after pharmacological isolation of eNMDAR-mediated responses. **(E)** Patch-clamp recordings from Ctrl and MHS hiPSC-neurons demonstrating pharmacological isolation of eNMDAR-mediated current and its inhibition by NitroSynapsin. Sample size listed in each panel. *^,#^*P* < 0.05, ****^,####^*P* < 0.0001 by ANOVA with post hoc Dunnett’s test for comparison with Ctrl (*) or with post hoc Sidak’s test between w/o and NitroSynapsin (^#^) for each genotype.

**Fig. S8.**
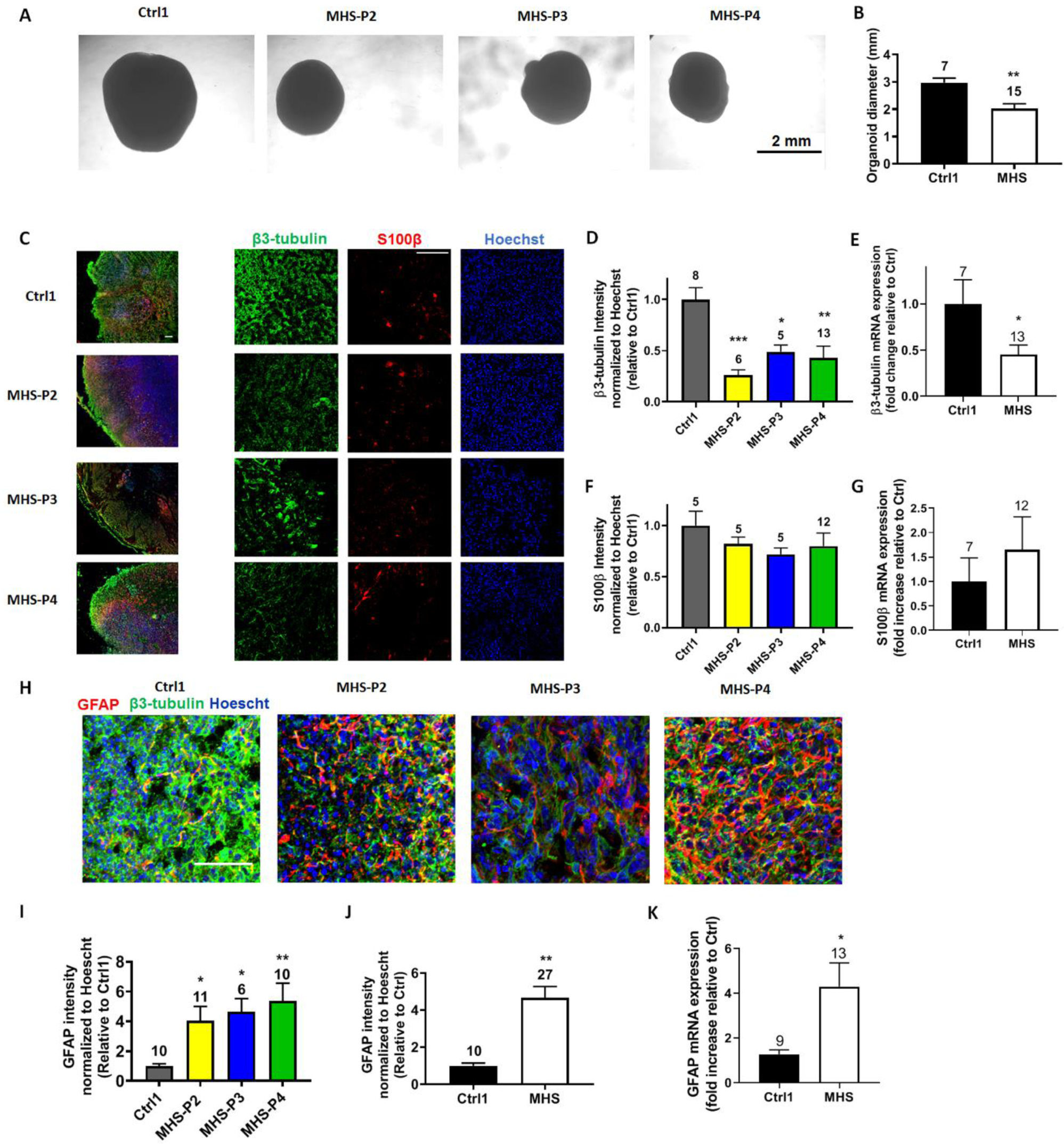
MHS hiPSC-derived cerebrocortical organoids generate fewer neurons. **(A)** Representative phase-contrast images of 2-month-old Ctrl and MHS organoids. Scale bar, 2 mm. (**B**) Quantification of the organoid diameter. (**C**) Representative images of β3-tubulin, S100β, and Hoechst for Ctrl and MHS hiPSC-neurons. *Left panel:* Low power images of whole organoids. *Right panel:* Higher power images of individual fluorescence channels. Scale bar, 100 µm. **(D)** Quantification of β3-tubulin fluorescence intensity for each MHS line normalized to Hoechst and relative to Ctrl. **(E)** β3-tubulin mRNA expression by qRT-PCR for grouped MHS hiPSC-neurons vs. Ctrl. **(F)** Quantification of S100β fluorescence intensity for each MHS line normalized to Hoechst and relative to Ctrl. **(G)** S100β mRNA expression by qRT-PCR in Ctrl vs. MHS as a group. (**H**) Representative images of GFAP, β3-tubulin, and Hoechst for Ctrl and MHS cerebral organoids. Scale bar, 50 µm. (**I**) Quantification of GFAP fluorescence intensity for MHS cerebral organoids normalized to Hoechst and relative to Ctrl. (**J**) Quantification of GFAP fluorescence intensity in Ctrl vs. MHS cerebral organoids as a group. **(K**) GFAP mRNA expression by qRT-PCR in Ctrl vs. MHS organoids as a group. Data are mean + SEM. Number of imaged fields (*n*) from 3 separate cerebral organoids (C and E) or number of organoids analyzed by qRT-PCR (C, E and F) are listed above bars. \**P* < 0.05; \*\**P* < 0.01; \*\*\**P* < 0.001 by ANOVA with Dunnett’s post hoc test for multiple comparisons or by unpaired Student’s t test for single comparisons.

**Fig. S9.**
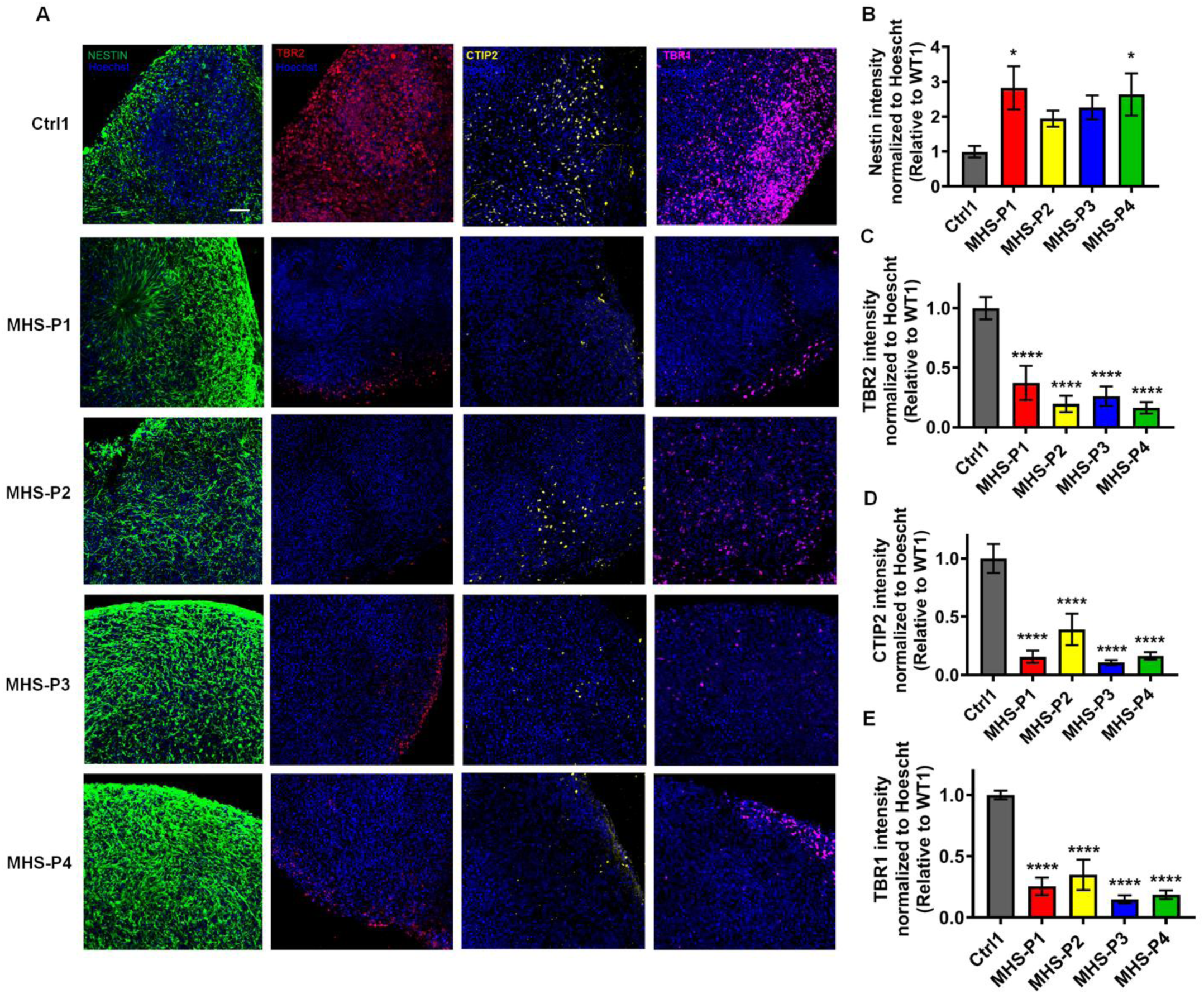
MHS hiPSC-derived cerebrocortical organoids display disrupted layering. (**A**) Representative images from left to right of Nestin, TBR2, CTIP2, TBR1, and Hoechst for Ctrl and MHS patients. Scale bar, 50 µm. **(B)** Quantification of Nestin fluorescence intensity for each MHS line relative to Ctrl. Fluorescence was normalized to Hoechst (labeling DNA in nucleus) to correct for cell number. **(C)** TBR2 fluorescence intensity for each MHS line normalized to Hoechst and relative to Ctrl. **(D)** CTIP2 fluorescence intensity for each MHS line normalized to Hoechst and relative to Ctrl. **(E)** TBR1 fluorescence intensity for each MHS line normalized to Hoechst and relative to Ctrl. Data are mean ± SEM. Sample size is 3-4 sections from 3 separate cerebral organoids for each genotype. \**P* < 0.05; \*\*\*\**P* < 0.0001 by ANOVA with Dunnett’s post hoc test.

**Fig. S10.**
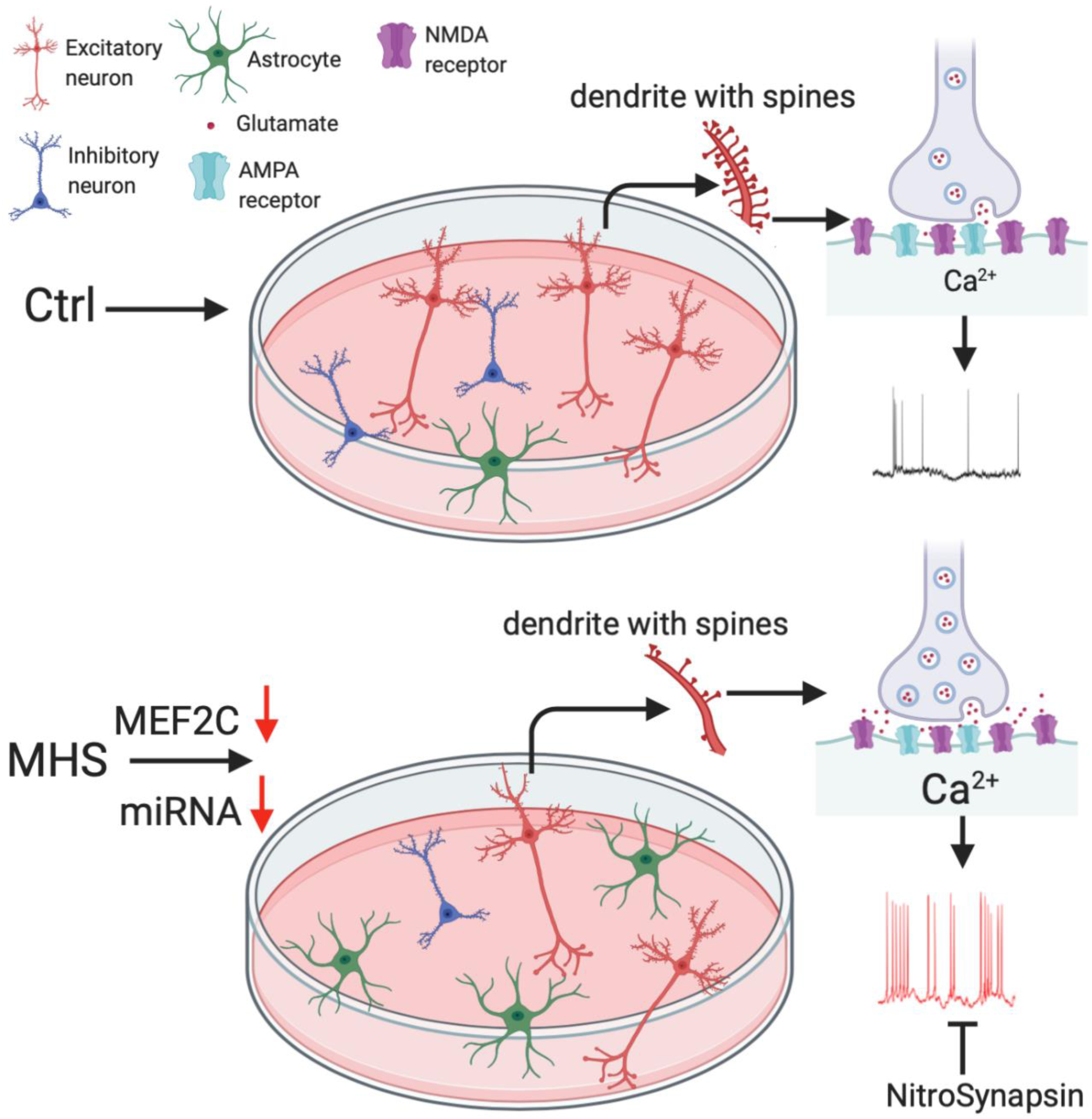
Aberrant gliogenesis and excitation in MEF2C autism patient hiPSC-neurons. Schematic diagram showing that MHS hiPSCs generate more astrocytes and fewer cerebrocortical neurons. Compared to control (Ctrl), the MHS neuronal population consists of fewer inhibitory neurons and excitatory neurons, but with more VGLUT1 vesicles, resulting in increased presynaptic glutamate release, increased postsynaptic intracellular Ca^2+^ levels, and hence increased excitability. The novel eNMDAR antagonist NitroSynapsin ameliorates this hyperactivity.

**Table S1.** Details of patients contributing fibroblasts for hiPSC generation.

| ID | Genotype | Age | Sex | Notable symptoms | hiPSC generated | Reference |
| --- | --- | --- | --- | --- | --- | --- |
| Ctrl1 | WT | neonatal | M | Isogenic control for MHS-P2 | Skin biopsy | 22 |
| Ctrl2 | WT | 18 y | M |  |  | 23 |
| Ctrl3 | WT | 8 y | F |  | Coriell |  |
| Ctrl4 | WT | 1 y | M |  | Skin biopsy |  |
| MHS-P1 | A106C (S36R) | 3 y | M | Averabal | Skin biopsy |  |
| MHS-P2 | Exon 1 del | neonatal | M | Isogenic to Ctrl1 by CRISPR/Cas9 |  |  |
| MHS-P3 | Exon 1 del | 6 y | M | Averbal, seizures | Skin biopsy | 16 |
| MHS-P4 | C52T (Q18X) | 4 y | F | Averbal, seizures | Skin biopsy |  |

**Table S2.** Electrophysiological properties of Ctrl and MHS hiPSC-derived cerebrocortical neurons.

| **Intrinsic properties** | **Ctrl1 (70)** | **Ctrl2 (25)** | **MHS-P1 (64)** | **MHS-P2 (40)** | **MHS-P3 (55)** | **MHS-P4 (60)** |
| --- | --- | --- | --- | --- | --- | --- |
| **Cell Capacitance (pF)** | 18.1±0.8 | 20.9±1.3 | 15.8±0.9 | 18.2±0.9 | 21.5±1.2 | 16.5±1.1 |
| **Sodium current density (pA/pF)** | 70.0±6.2 | 59.4±10.2 | 65.0±8.5 | 85.6±10.8 | 79.6±7.7 | 64.1±4.0 |
| **Potassium current density (pA/pF)** | 96.0±6.7 | 100.5±7.4 | 120.1±13.5 | 113.2±20.4 | 93.9±9.0 | 117.2±10.0 |
| **Resting Membrane Potential (mV)** | -30.5±1.1 | -30.1±1.7 | -28.4±1.3 | -31.6±1.3 | -32.9±1.6 | -29.1±1.0 |
| **Action Potential (AP) height (mV)** | 91.9±2.4 | 87.2±2.6 | 89.7±2.4 | 97.6±2.0 | 96.3±1.8 | 91.5±1.9 |
| **AP half-width (ms)** | 8.5±0.5 | 9.4±0.8 | 8.6±0.6 | 7.3±0.6 | 7.0±0.5 | 7.5±0.4 |
| **AP threshold (mV)** | -28.4±0.5 | -29.9±1.0 | -27.9±0.6 | -27.1±0.8 | -28.9±0.8 | -26.8±0.6 |

**Table S3.** ChIP-seq targets of transcription factor MEF2 (see separate excel file).

**Movie S1**. Ctrl hiPSC-derived cerebrocortical neuron and astrocyte cultures. Fluo-4 measurement of intracellular Ca^2+^ showing low basal endogenous signaling activity.

**Movie S2**. MHS hiPSC-derived cerebrocortical neuron and astrocyte cultures. Fluo-4 measurement of intracellular Ca^2+^ showing high synchronous burst-like basal endogenous signaling activity.

**Movie S3**. Effect of NitroSynapsin on MHS hiPSC-derived cerebrocortical neuron and astrocyte cultures. Fluo-4 measurement of intracellular Ca^2+^ showing increased excitability, which was normalized after application of 10 µM NitroSynapsin.

